# EGFR-initiated endocytosis of Wnt9a and Fzd9b is required for β-catenin signaling

**DOI:** 10.1101/2022.09.25.509379

**Authors:** Nicole Nguyen, Kelsey A. Carpenter, Kate E. Thurlow, Emily Mu, Carla Gilliland, Stephanie Grainger

## Abstract

Cell to cell communication through secreted ligands like those encoded by the *Wnt* gene family is critical for development and homeostasis during organismal life. One of the bottlenecks in the Wnt field has been identifying specific ligand/receptor pairings and decoding the mechanisms for their downstream signals. We previously discovered that the Wnt9a ligand signals through the cell surface receptors Fzd9b, LRP5/6 and EGFR to promote early proliferation of hematopoietic stem cells during development. We used this exquisitely specific ligand/receptor complex as a platform to determine if Wnt9a requires endocytosis for signaling. Using fluorescently labeled, biologically active Wnt9a and Fzd9b fusion proteins, we demonstrate here that the Wnt9a receptor complex is rapidly endocytosed within one minute of contact with Fzd9b. Following this, the Wnt9a/Fzd9b complex is trafficked through the cell to early and late endosomes, lysosomes, and the endoplasmic reticulum; it is also recycled back to the membrane. Using small molecule inhibitors, genetic and siRNA approaches, we identified that mechanistically this endocytosis requires EGFR-mediated phosphorylation of the Fzd9b tail, followed by endocytosis through a caveolin and EPS15 dependent pathway. Specific modes of endocytosis and trafficking may represent one of the ways in which Wnt/Fzd specificity is established, since other Wnt ligands do not require endocytosis for activity.

## Introduction

Intercellular signaling is essential for the normal development and homeostasis of organismal life. Secreted: secreted signaling molecules direct neighboring cells to adopt different cellular fates, maintain pluripotency, and/or stimulate cellular behaviors. One vital class of cell signaling molecules are the highly conserved Wnt proteins, which elicit cellular signals that orchestrate a plethora of key biological processes, including cellular symmetry, proliferation, tissue polarity and stem cell maintenance^1, 2^.

Wnt signaling is generally separated into “canonical” and “non-canonical” pathways^1, 2^; this study focuses on the canonical pathway, which is dependent upon the nuclear translocation of the transcriptional activator β−catenin. At the cell membrane, Wnt signals are generally initiated when one of many Wnt ligands interacts with one of several cell surface receptors. The *Frizzled (Fzd)* gene family encodes a class of seven-pass transmembrane receptors that can bind Wnt ligands through a C-terminal extracellular cysteine rich domain (CRD)^3^, as well as co-receptors such as those from the *lipoprotein-rich protein (LRP)* gene family. A large number of Wnt ligands (≥19) and Fzd receptors (≥10) are encoded in vertebrate genomes; however, the specificity of signaling interactions has only begun to be elucidated^4–7^.

We identified an exquisitely specific pairing between Wnt9a and Fzd9b in zebrafish hematopoietic stem and progenitor cell (HSPC) development^8, 9^. This pairing is also conserved in human cells^8–10^. Surprisingly, Epidermal Growth Factor Receptor (EGFR) is a required co-factor for this signal^8^; however, we do not yet understand the molecular mechanisms downstream of this requirement.

One mechanism for differentially deciphering and propagating cell signals is through endocytosis and sorting. Generally, cargoes at the cell membrane are endocytosed through different pathways including those mediated by clathrin or caveolin, among others^11, 12^. These nascent endosomes will fuse together and recruit endosomal proteins to become early endosomes. Following this, cargoes are either (1) sorted to the late endososome/multi-vesicular body (MVB), or (2) exported to recycling endosomes to be transported back to the plasma membrane. From the late endosome/MVB, cargoes can be sorted to the lysosome for degradation, or to the Golgi apparatus for reuse at the membrane^13, 14^. There is now a substantial body of evidence supporting a requirement for different endocytosis pathways for signal activation, sustaining a signal, or for differential signaling for many receptors^15–25^. There are conflicting studies on the requirement(s) for endocytosis in Wnt signaling, possibly due to cell- and receptor/ligand-dependent contexts for downstream signals^26–38^.

We developed fluorescently tagged Wnt and Fzd molecules expressed in CHO and HEK293T cells, respectively. These tagged proteins are expressed, properly trafficked, and biologically active. Using this platform, we observed that Wnt9a/Fzd9b complexes are internalized within one minute of treatment with Wnt9a. Interestingly, small molecule inhibition of endosome scission abrogates the Wnt9a/Fzd9b-dependent β-catenin signal, while inhibition of lysosomal degradation increases this signal, suggesting that Wnt9a/Fzd9b signals from endosomes. Mechanistically, this internalization event requires EGFR tyrosine phosphorylation of Fzd9b; mutation of Fzd9b to eliminate the EGFR phosphorylation site that we previously identified^8^ also eliminates internalization of the Wnt9a/Fzd9b complex. Internalization of the receptor complex requires caveolin-mediated endocytosis and the scaffolding protein EGFR protein substrate 15 (EPS15)^39–43^. We also observed that Wnt9a/Fzd9b signaling is dependent on EPS15. Finally, we found that EGFR tyrosine phosphorylation sites on EPS15 and Fzd9b as well as the previously described EH (Eps homology) domain of EPS15 are all required for Wnt9a/Fzd9b signaling^41, 44^.

## Results

### Fluorescently tagged Wnt9a and Fzd9b proteins maintain signaling activity

There are conflicting reports on the importance of endocytosis for Wnt signaling for various Wnt/Fzd pairings^26–38^. To investigate how Fzd9b responds to its cognate ligand, Wnt9a, we generated constructs expressing either Fzd9b or Fzd9b lacking the ligand-binding cysteine rich domain (CRD), fused to a 5(GS) linker sequence and either the red fluorescent protein mKate2, or a V5 tag, (hereafter Fzd9b-mKate, Fzd9b-V5, ΔCRD Fzd9b-mKate or ΔCRD Fzd9b-V5, Fig. 1A, Fig. S1A). The mKate fusion proteins were stably expressed under regulatory control of a cytomegalovirus (CMV) promoter in HEK293 cells, and were detected by immunoblotting at the anticipated sizes using either our previously described Fzd9b antibody^8^ or an mKate antibody (Fig. 1B). Using confocal microscopy and immunofluorescence, we saw strong overlap between the N-terminal Fzd9b antibody staining, and the C-terminal mKate fluorescent reporter (Fig. 1C). Results were the same with a V5 antibody (Fig. S1B), indicating that mKate or V5 can be used as a surrogate reporter of Fzd9b localization in the cell. Wnt signaling through β-catenin can be measured using a luciferase-based reporter assay called Super TOP (TCF-optimal promoter) Flash (STF)^45^. We found that tagging Fzd9b with either mKate or V5 did not impair its signaling ability, and that as anticipated, ΔCRD Fzd9b-mKate did not display any signaling ability (Fig. 1D, Fig S1C). Finally, we tested that these fusion proteins were correctly trafficked to the cell membrane using non-permeabilized immunofluorescence with our extracellular Fzd9b antibody and found that Fzd9b fusion proteins were correctly trafficked to the cell surface (Fig. S1D-E).

**Figure 1:**
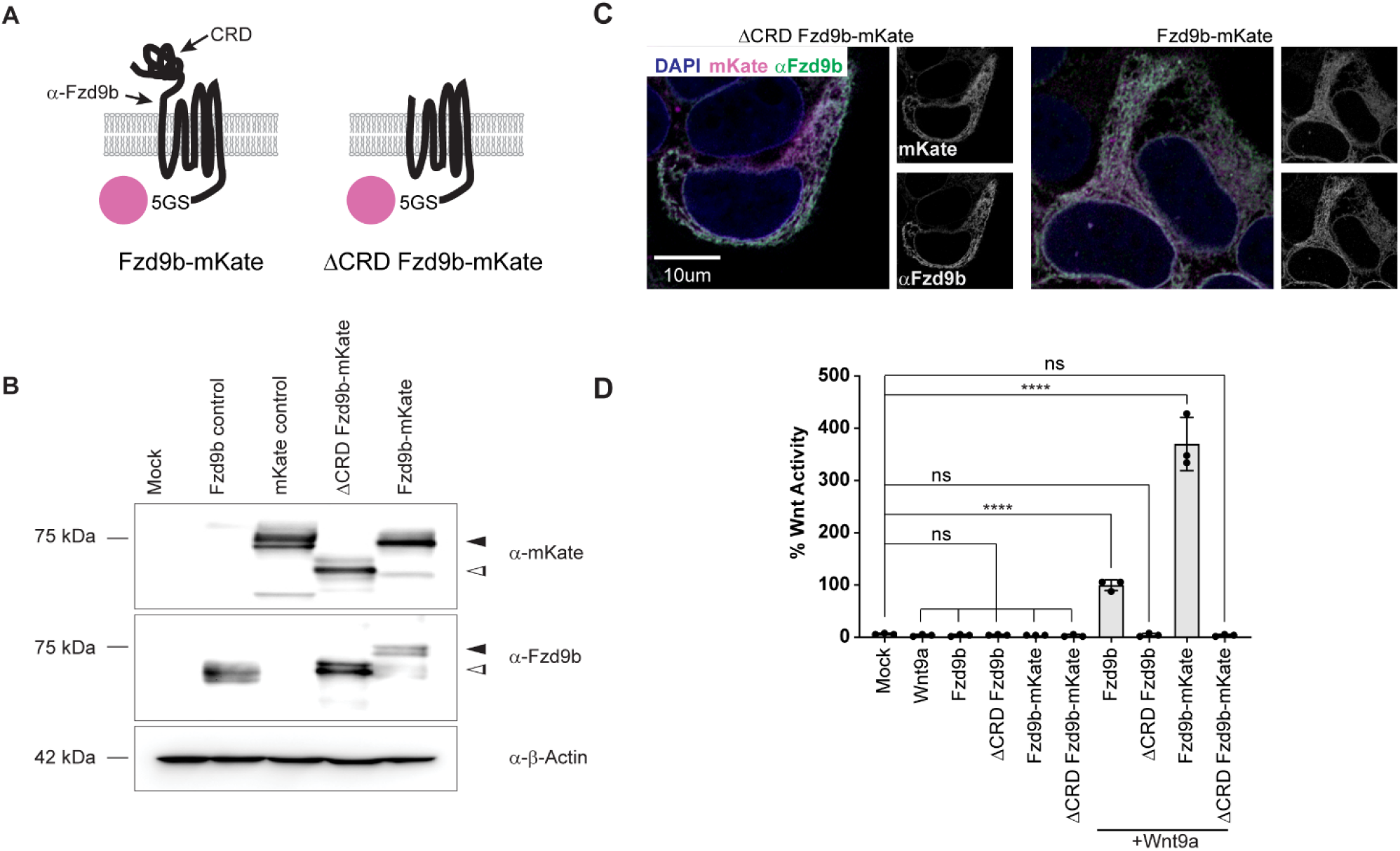
Fluorescently tagged Fzd receptors maintain signaling capacity. **A.** Schematic of Fzd9b-mKate and ΔCRD Fzd9b-mKate constructs. **B.** Confocal z-stacks of transgenic lines as indicated. Magenta is mKate-fused Fzds, blue is DAPI (nuclei), green is α-Fzd9b. **C**. Immunoblots for mKate, Fzd9b and β-Actin from transgenic HEK293T cell lines stably expressing Fzd constructs as shown. **D.** STF assays for Fzd fusion proteins co-transfected with WT Wnt9a of GFP mock control plasmid. n.s. not significant, ****P<0.0001 by ANOVA with Tukey post-hoc comparison compared to mock.

### A Gamillus-tagged Wnt9a protein retains biological activity

There are only a handful of previously successful instances of generating functional Wnt proteins tagged with fluorescent proteins^46–51^, likely due to their hydrophobic nature, which is a requirement for their signaling activity. We began by generating versions of Wnt9a tagged with a 5(GS) linker fused to either FLAG or Green Fluorescent Protein (GFP) at the N-terminus between L58 and C59, or at the C-terminus, just before the stop codon (hereafter FLAG-Wnt9a, GFP-Wnt9a or Wnt9a-GFP, Fig. S2A), under regulatory control of a doxycycline inducible promoter system. We generated stable transgenic cell lines in CHO cells and found that all three of these fusion proteins were expressed and secreted; however, both GFP-tagged Wnt9a proteins showed some degree of presumed cleavage (Fig. S2B). Conditioned medium from FLAG-Wnt9a or Wnt9a-GFP were able to induce STF activity with Fzd9b, suggesting that these tagged versions of Wnt9a are biologically active (Fig. S2C). On the other hand, GFP-Wnt9a did not induce STF reporter activity (Fig. S2C), indicating that the N-terminal GFP fusion protein is not active in STF assays.

As cargo is endocytosed, it is acidified through the endosome-lysosome pathway, which abolishes the fluorescent output of some fluorophores like GFP, but not Gamillus, another fluorescent protein in the green range^52^. To monitor Wnt9a as it is endocytosed, we derived C-terminally tagged Wnt9a constructs fused with either 5(GS), 3(GAS) or GTG-2(SGGGG)SGGS linker sequences and Gamillus (Fig. 2A). Although all three of these fusions were expressed and secreted from CHO cells, we observed putative cleavage of the 5(GS) and 3(GAS) fusion proteins (Fig. 2B). The Wnt9a-GTG-2(SGGGG)SGGS-Gamillus (hereafter Wnt9a-Gam) remained intact (Fig. 2B), and so we chose to pursue this as our reporter of Wnt9a localization. Immunofluorescent staining of Wnt9a-Gam CHO cells showed colocalization of the Wnt9a antibody that we previously derived^10^ and the Gamillus fluorescent signal (Fig. 2C). We also observed overlapping labeling of FLAG and Wnt9a in FLAG-Wnt9a CHO cells (Fig. S2D). Treating Fzd9b-mKate STF reporter cells with Wnt9a-Gam conditioned medium showed nearly wild-type levels of STF activity, indicating that this fusion protein is biologically active (Fig. 2D). Taken altogether, these data indicate that Wnt9a-Gam can be used to visualize Wnt9a.

**Figure 2:**
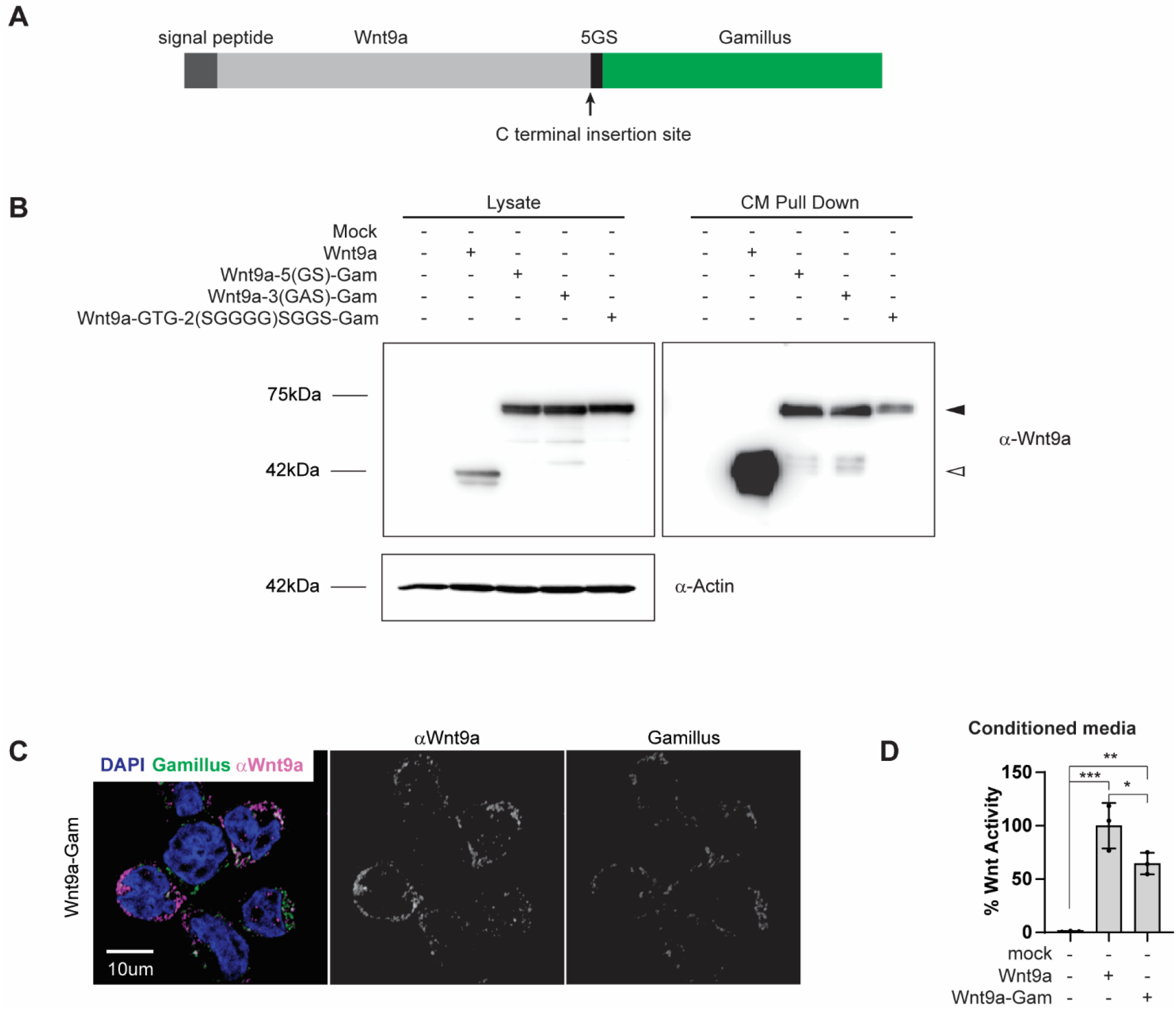
A Wnt9a fusion protein maintains signaling. **A.** Schematic of Wnt9a-Gam construct. **B.** Immunoblots for Wnt9a and β-Actin; cell lysates and conditioned media (CM) before and after Wnt9a pulldown are shown for transgenic CHO cell lines as indicated. Black arrowhead indicated anticipated size for Wnt9a fused to Gam; white arrowhead indicates anticipated size of untagged Wnt9a. **C.** Confocal z-stacks of immunofluorescence on transgenic lines as shown. Magenta is α-Wnt9a, green is Gamillus, blue is DAPI. **D.** STF assays conducted in cells stably expressing Fzd9b-mKate and STF and induced with CM from transgenic CHO lines expressing Wnt9a or Wnt9a-Gam. n.s. not significant and *P<0.05, **P<0.01, ***P<0.001 by ANOVA with Tukey post-hoc comparison.

### The Wnt9a/Fzd9b complex is rapidly endocytosed

Endocytosis is a common mechanism of signal regulation for a variety of cell signaling cascades, and some receptor complexes require endocytosis to initiate different downstream signals^53–58^. To assess if the Wnt9a/Fzd9b complex undergoes endocytosis, we treated Fzd9b-mKate or ΔCRD Fzd9b-mKate cells with conditioned medium collected from either naïve CHO (mock CM) or Wnt9a-Gam CHO (Wnt9a-Gam CM) cells and fixed the cells after several timepoints for analysis by confocal microscopy. A small slice through the center of the Fzd9b-mKate cells indicates that, in response to Wnt9a-Gam treatment, there was rapid internalization (as early as 15 seconds and most robustly at 1 minute) of co-localized Wnt9a and Fzd9b (Fig. 3A-B), a finding we recapitulated using Wnt9a-GFP (Fig. S3A). We occasionally detected very small and faint Wnt9a/Fzd9b complexes in later timepoints such as 6 hours post treatment (Fig. 3A-B). On the other hand, although ΔCRD Fzd9b-mKate cells had multiple Fzd9b puncta on the inside of the cell, none of these we co-localized with Wnt9a (Fig. 3A), indicating that the CRD is required for Wnt9a internalization.

**Figure 3:**
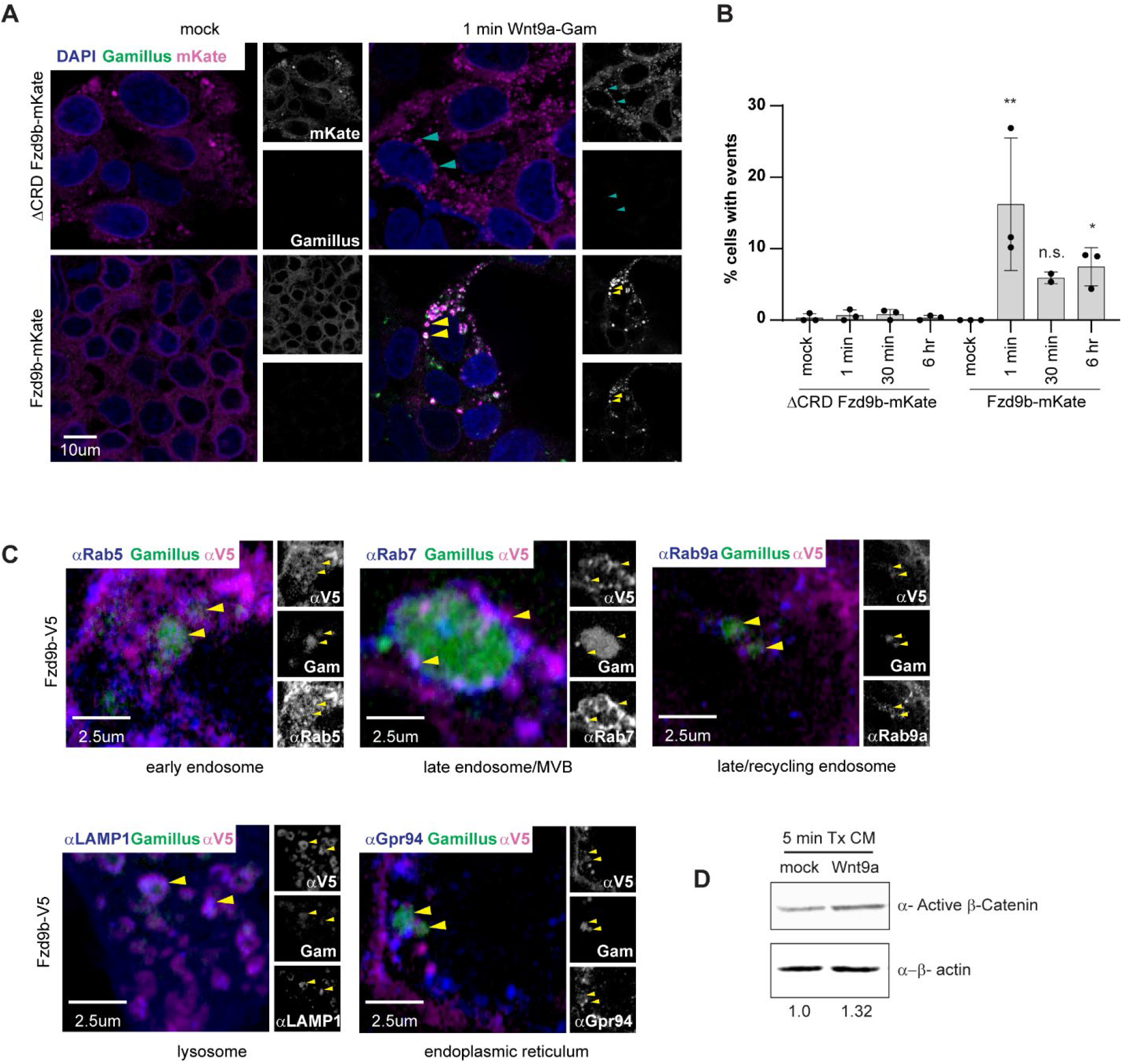
The Wnt9a/Fzd9b complex is rapidly endocytosed. **A.** Confocal z-stacks of ΔCRD Fzd9b-mKate or Fzd9b-mKate cell lines treated with either mock CM or Wnt9a-Gam CM after 1 minute. Yellow arrows indicate Wnt9a/Fzd9b complexes; cyan arrows indicate Fzd9b without Wnt9a. **B.** Quantification of cells with internalized Wnt9a-Gam from **A**. **C.** Confocal z-stacks of immunofluorescence on Fzd9b-V5 cells treated for one minute with Wnt9a-Gam. Magenta is α-V5 (Fzd9b), green is Gamillus, blue is endocytosis markers as shown. **D.** Immunoblot for active β-catenin (using an antibody targeted to non-phosphorylated β-catenin) in Fzd9b-mKate cells treated with mock CM or Wnt9a-Gam CM for 5 minutes. n.s. not significant and *P<0.05, **P<0.01 by ANOVA with Tukey post-hoc comparison.

Passage through the endosomal trafficking pathway leads to the interaction of cargoes with various Rab GTPases that mark subcellular structures throughout the cell. By co-staining Fzd9b-V5 cells treated with Wnt9a-Gam, we were able to detect Wnt9a/Fzd9b complexes in the early endosome (Rab5), late endosome/multi-vesicular body (Rab7), late/recycling endosome (Rab9a), endoplasmic reticulum (Gpr94), and lysosome (LAMP1) (Fig. 3C), indicating that Wnt9a/Fzd9b complexes are rapidly endocytosed and trafficked throughout the cell.

There are many mechanisms through which cells uptake cargoes, including clathrin-and caveolin-mediated endocytosis^59^. To determine if endocytosis was required for Wnt9a/Fzd9b signaling, we used the small molecule inhibitors, PitStop2 and Dynasore, which inhibit both clathrin-dependent and clathrin-independent endocytosis^60, 61^. We found that these inhibited Wnt9a/Fzd9b-dependent STF reporter activity in a dosage and Wnt-dependent manner (Fig. S3B-C), suggesting that Wnt9a/Fzd9b endocytosis is required for Wnt signaling. Once cargoes are endocytosed, they are sorted to various intracellular destinations, including to the lysosome (Fig. 3C). Using Bafilomycin A1, an inhibitor of the V-ATPases critical for lysosomal degradation, we observed a dosage-dependent increase in Wnt9a/Fzd9b STF activity (Fig. S3D). These data were also supported by data where knocking down two of the V-ATPases required for lysosomal degradation, using siRNA in Fzd9b STF cells, leads to an increase in Wnt9a/Fzd9b-driven STF activity (Fig. S3E). Taken together with the observation that internalization is required for signaling activity, these results suggest that the Wnt9a/Fzd9b receptor complex signals from endosomes.

In BCD signaling, once a Wnt ligand interacts with its cognate receptor, Fzd, the β-catenin destruction complex is dissociated, allowing non-phosphorylated (active) β-catenin to enter the nucleus and activate target gene expression^62, 63^. The observation of rapid endocytosis of the Wnt9a/Fzd9b complex, paired with the requirement for endocytosis in initiating signaling suggested that there should be very early activation of β-catenin in response to Wnt9a, which we observed within 5 minutes of Wnt9a treatment (Fig. 3D). Altogether, these results indicate that Wnt9a/Fzd9b are rapidly endocytosed, and that this is a prerequisite for β-catenin dependent signaling.

### Caveolin-dependent endocytosis is required for Wnt9a/Fzd9b signaling

We had previously observed that key components of the AP-2 complex, which is essential for clathrin-mediated endocytosis, are recruited to Fzd9b in response to Wnt9a^8^. To test the requirement of clathrin-mediated endocytosis in Wnt9a/Fzd9b signaling, we used an siRNA targeting the *clathrin heavy chain* (*CLTC*) in Fzd9b STF cells, and tested STF activity in response to Wnt9a. While we observed approximately 92% knockdown of CLTC protein (Fig. S4A), there was no effect on Wnt9a/Fzd9b STF activity (Fig. S4B). This was not due to residual CLTC protein, since the prototypical clathrin cargo Transferrin was not endocytosed under knockdown conditions (Fig. S4C), altogether indicating that clathrin heavy chain is not required for Wnt9a/Fzd9b signaling.

To determine if caveolin-mediated endocytosis may mediate Wnt9a/Fzd9b endocytosis, we used an siRNA targeting *Caveolin* (*CAV1)*, an essential component of caveolin pits, and found that in the context of undetectable amounts of CAV1 protein (Fig. 4A), the Wnt9a/Fzd9b signal was severely compromised (Fig. 4B). To determine if CAV1 impacts Wnt9a/Fzd9b endocytosis, we treated Fzd9b-mKate cells with either mock CM or Wnt9a-Gam CM in the context of either control or *CAV1* siRNA sequences and examined cells at 1 minute post CM treatment. Using this approach, we found that in the absence of CAV1, Wnt9a/Fzd9b endocytosis was severely compromised (Fig. 4C-D), indicating that the Wnt9a/Fzd9b complex is internalized through caveolin-mediated endocytosis.

**Figure 4:**
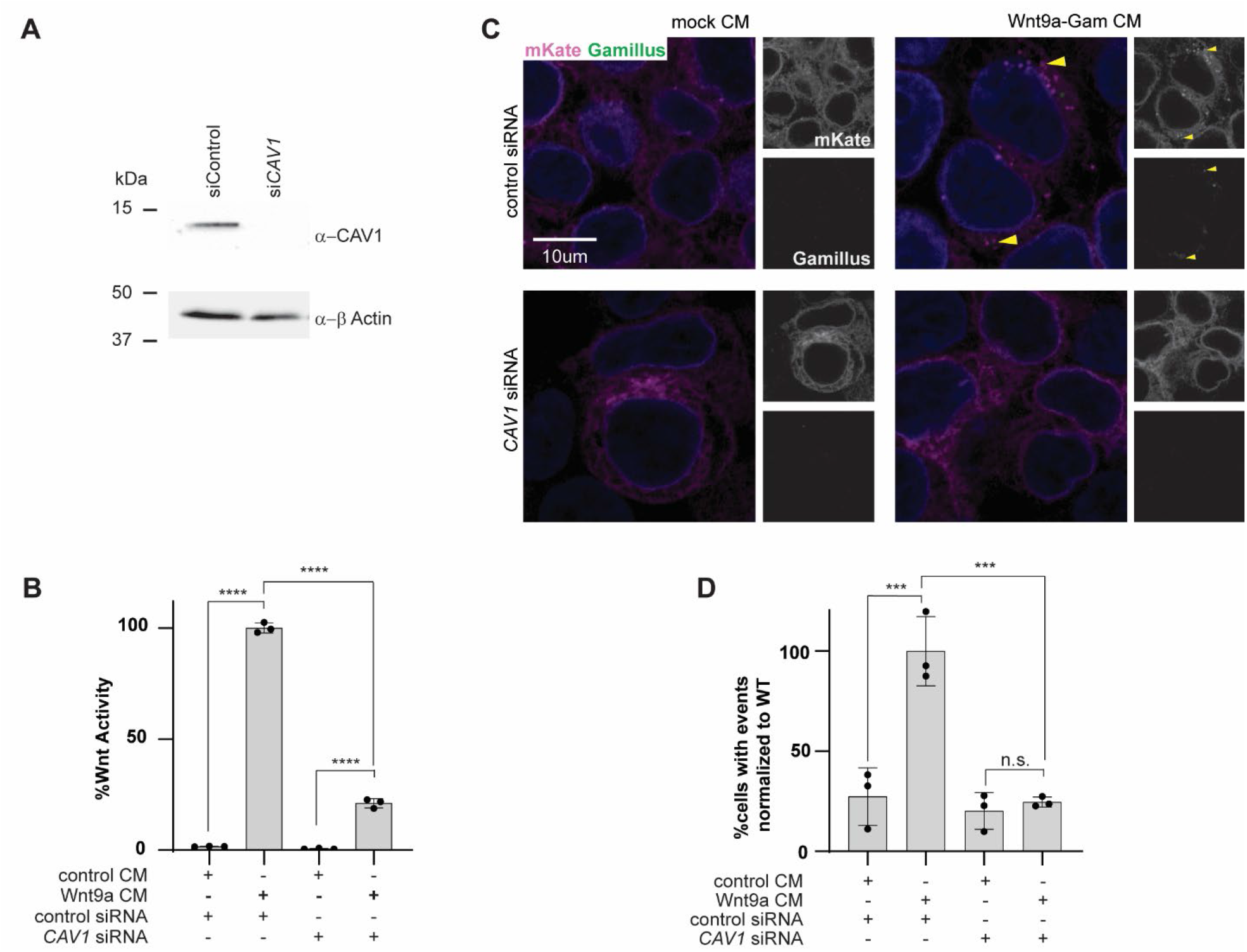
Caveolin-dependent endocytosis is required for Wnt9a/Fzd9b signaling. **A.** Immunoblot of Fzd9b-mKate cells treated with either control or *CAV1* siRNAs. **B.** STF assays conducted in cells stably expressing Fzd9b-mKate and STF, treated with either control or *CAV1* siRNA, and induced with either mock or Wnt9a CM. **C.** Confocal z-stacks of Fzd9b-mKate cells treated with either control or *CAV1* siRNA, induced with either mock CM or Wnt9a-Gam CM, and fixed after one minute. Magenta is mKate, green is Gamillus, blue is DAPI, quantified in **D.** n.s. not significant and ***P<0.001, ****P<0.0001 by ANOVA with Tukey post-hoc comparison.

### Wnt9a/Fzd9b endocytosis does not require LRP

We next sought to identify the mechanism through which endocytosis of Wnt9a/Fzd9b is initiated at the membrane. The co-receptors Low-density lipoprotein receptor-related protein 5 and 6 (LRP5 and LRP6) are required for the Wnt9a/Fzd9b signal^8^, and it is thought that dissociation of the destruction complex is initiated when the key destruction complex scaffold protein Axin is recruited to LRP5/6^64–66^. To determine if LRP5 and LRP6 are required for Wnt9a/Fzd9b endocytosis, we generated LRP5/6 double knockout (DKO) HEK293 STF cells using CRISPR/Cas9 in our previously established LRP6 knockout cells^8^. We isolated a clone with 10 bp deletion and 1 bp insertion alleles, which were both predicted to cause frameshift mutations and loss of LRP5 function (Fig. S5A); we did not detect any LRP5 or LRP6 protein in these LRP5/6 DKO cells (Fig. S5B). As expected, these cells maintained their ability to respond to the Wnt agonist CHIR, which operates downstream of the receptor complex, but did not respond to either the prototypical ligand Wnt3a, or to Wnt9a, or Wnt9a-Gam in the presence of Fzd9b (Fig. S5C).

To determine if Wnt9a/Fzd9b complex endocytosis relies upon LRP5/6, we generated LRP5/6 DKO cells which transgenically expressed Fzd9b-mKate and treated these with either mock CM or Wnt9a-Gam CM and examined cells for Wnt9a/Fzd9b endocytosis after 1 minute. We found that in the absence of LRP5/6, Wnt9a/Fzd9b endocytosis was unaffected (Fig. S5D), indicating that while LRP5/6 are required for β-catenin signaling, they are not required for Wnt9a/Fzd9b endocytosis.

### Wnt9a/Fzd9b endocytosis requires EGFR-mediated phosphorylation of the Fzd9b tail

We previously determined that EGFR is an obligate co-factor for the Wnt9a/Fzd9b signal: in response to Wnt9a, EGFR is required to phosphorylate tyrosine 556 (Y556) on the Fzd9b cytoplasmic tail for signal initiation, independent of the canonical EGFR signaling pathway^8^. To study the function of EGFR in Wnt9a/Fzd9b endocytosis, we generated EGFR knockout (KO) HEK293 cells using CRISPR/Cas9. We identified a clone with several early mutations in the EGFR coding sequence (Fig. S5E). These cells had undetectable levels of EGFR protein (Fig. S5F) and lacked tyrosine phosphorylation activity (Fig. S5G). EGFR is a specific co-factor for the Wnt9a/Fzd9b signal and as expected, EGFR KO cells were able to respond to Wnt3a and CHIR, but not to Wnt9a (Fig. S5H).

To determine if EGFR is required for Wnt9a/Fzd9b endocytosis, we generated transgenic Fzd9b-mKate cells in the EGFR KO background, and treated these or control EGFR WT cells with either mock CM or Wnt9a-Gam CM. In the absence of EGFR, Wnt9a/Fzd9b endocytosis was nearly completely absent (Fig. 5A-B). To test if the enzymatic function of EGFR is required for this endocytic event, we used EGFR WT Fzd9b-mKate cells treated with Wnt9a-Gam or mock CM in the presence of the EGFR tyrosine kinase inhibitor AG-1478 or vehicle and found that AG-1478 also compromised Wnt9a/Fzd9b endocytosis (Fig. 5C-D). We next compared the capacity for Fzd9b Y556F, which lacks the ability to be phosphorylated by EGFR in response to Wnt9a^8^, to be endocytosed in response to Wnt9a. We found that in the absence of this key tyrosine phosphorylation site, Wnt9a/Fzd9b endocytosis was severely hampered (Fig. 5E-F). Altogether, these results indicate that EGFR-mediated phosphorylation of the Fzd9b tail at Y556 is required for Wnt9a/Fzd9b endocytosis.

**Figure 5:**
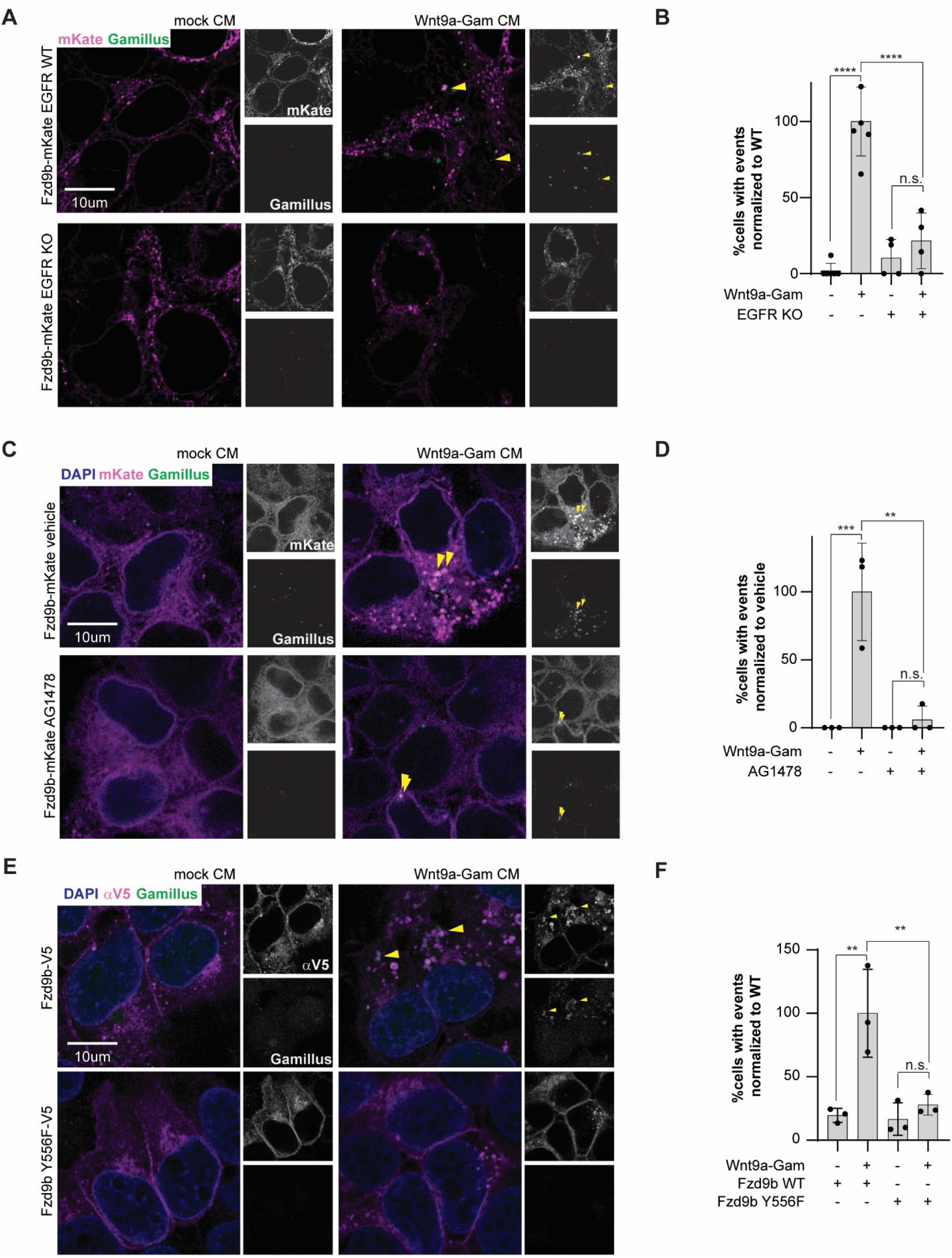
Wnt9a/Fzd9b trafficking requires EGFR-initiated phosphorylation of the Fzd9b tail at Y556. **A.** Confocal z-stacks of Fzd9b-mKate WT or EGFR KO cells, induced with either mock CM or Wnt9a-Gam CM, and fixed after one minute. Magenta is mKate, green is Gamillus, blue is DAPI, quantified in **B. C.** Confocal z-stacks of Fzd9b-mKate WT cells, treated with either vehicle or 2.5μM AG1478, and induced with either mock CM or Wnt9a-Gam CM, and fixed after one minute. Magenta is mKate, green is Gamillus, blue is DAPI, quantified in **D. E.** Confocal z-stacks of Fzd9b WT-mKate or Fzd9b Y556F-mKate cells, induced with either mock CM or Wnt9a-Gam CM, and fixed after one minute. Magenta is mKate, green is Gamillus, blue is DAPI, quantified in **F.** n.s. not significant and **P<0.01, ***P<0.001, ****P<0.0001 by ANOVA with Tukey post-hoc comparison.

### EPS15 is required for Wnt9a/Fzd9b endocytosis and signaling

During EGFR signaling events, the EGFR receptor can be endocytosed through multiple mechanisms, and these lead to distinct transcriptional responses^53, 54, 57, 58^. One of the key mediators of EGFR endocytosis is the adaptor protein EPS15, which also has roles in endocytosis of other receptors^39, 41, 67–69^. To identify proteins that are localized to the Fzd9b tail, we had previously conducted an APEX2-mediated proximity ligation screen^8^, where we found that EPS15 is enriched to nearly the same degree as EGFR (Fig. S6A). We used Wnt9a treatment of Fzd9b STF cells in the context of either control or *EPS15* siRNA sequences and found that under conditions of undetectable EPS15 protein (Fig. S6B), Wnt9a/Fzd9b signaling is severely compromised (Fig. 6A).

**Figure 6:**
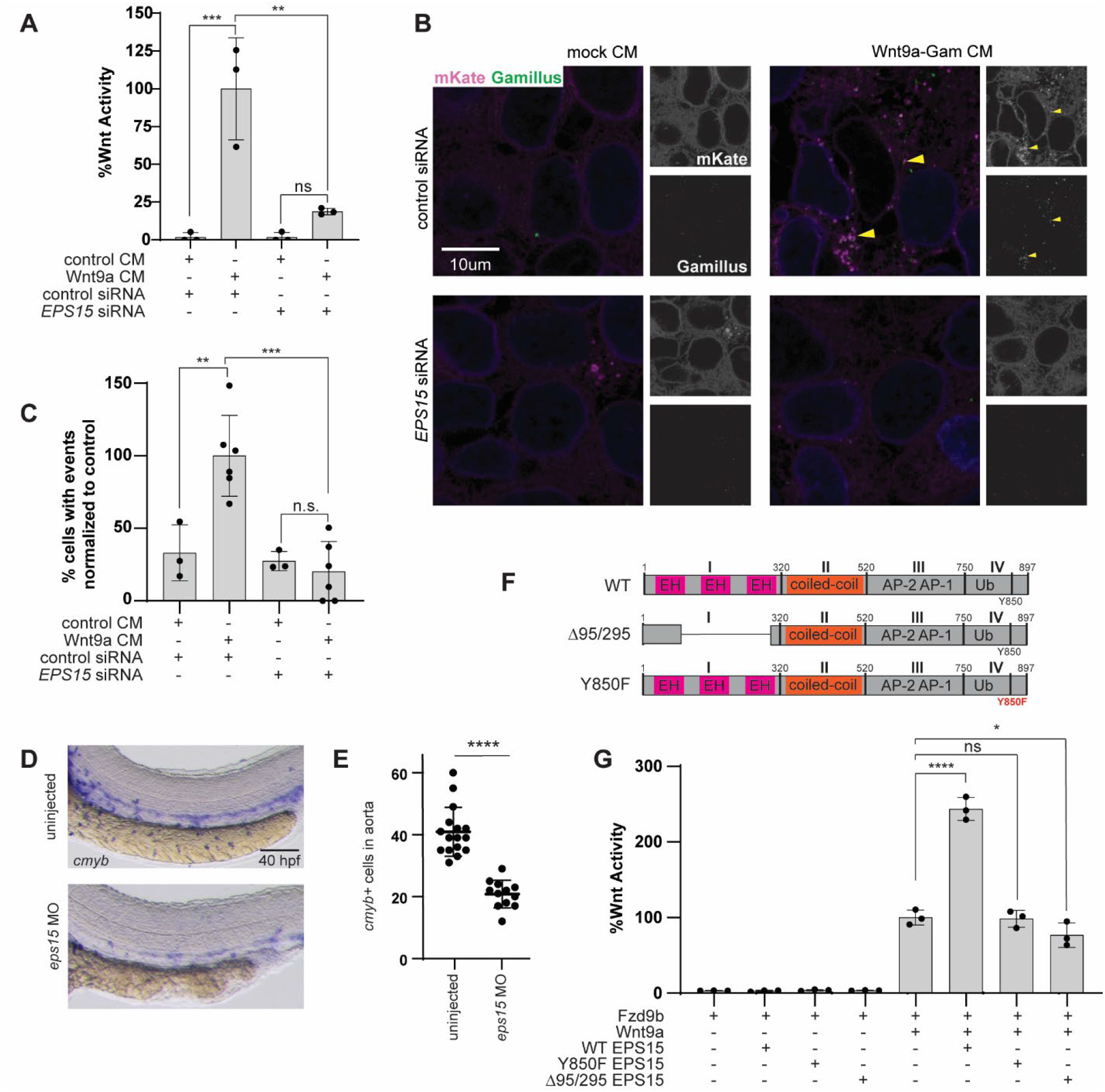
EPS15 is required for Wnt9a/Fzd9b endocytosis and signaling. **A.** STF assays conducted in cells stably expressing Fzd9b-mKate and STF, treated with either control or *EPS15* siRNA, and induced with either mock or Wnt9a CM. **B.** Confocal z-stacks of Fzd9b-mKate cells, treated with either control or *EPS15* siRNA, and induced with either mock CM or Wnt9a-Gam CM, and fixed after one minute. Magenta is mKate, green is Gamillus, blue is DAPI, quantified in **C. D.** Zebrafish at 40 hpf stained for hematopoietic stem and progenitor cell maker *cmyb* by *in situ* hybridization, quantified in **E**. **F.** EPS15 construct schematics. **G.** STF assays conducted in cells stably expressing Fzd9b-mKate and STF, transfected with EPS15 constructs, and induced with either mock or Wnt9a CM. n.s. not significant and *P<0.05, **P<0.01, ***P<0.001, ****P<0.0001 by ANOVA with Tukey post-hoc comparison.

To determine if EPS15 plays a role in Wnt9a/Fzd9b receptor complex internalization, we examined Wnt9a/Fzd9b internalization after 1 minute of Wnt9a-Gam treatment in the context of control or EPS15 siRNA sequences and found a loss of endocytosis of Wnt9a/Fzd9b without EPS15 (Fig. 6B-C), indicating that EPS15 plays a role in Wnt9a/Fzd9b endocytosis.

We have previously shown that the Wnt9a/Fzd9b signal is required *in vivo* for HSPC proliferation; defects in this process can be visualized using *in situ* hybridization for the HSPC marker *cmyb* at 40 hours post fertilization (hpf) in zebrafish^8, 9^. To identify if EPS15 plays a role in HSPC proliferation, we injected an antisense morpholino (MO) targeting EPS15 into wild-type zebrafish zygotes and analyzed the *cmyb*^+^ cells in the floor of the dorsal aorta at 40 hpf, where we found a loss of *cmyb*^+^ cells in EPS15 MO animals (Fig. 6D). These results indicate that EPS15 is required for HSPC development and support its function in the Wnt9a/Fzd9b pathway.

EPS15 has been subdivided into four main domains (I-IV, Fig. 6F). A deletion of residues 95-295 (Δ95/295, most of domain I) has been reported to be required for transferrin receptor and EGFR endocytosis^44^; the Y850 residue has been reported as a substrate for EGFR required for endocytosis^41^. To test if these EPS15 mutants impacted on the Wnt9a/Fzd9b STF signal, we transfected either WT, Y850F or Δ95/295 into our Wnt9a/Fzd9b STF reporter assay. While we observed approximately 2.5x enhanced Wnt9a/Fzd9b activity in the context of WT EPS15, the signal was not enhanced by either Y850F or Δ95/295 EPS15 (Fig. 6G); we observed very slight dominant-negative activity with Δ95/295 EPS15 (Fig. 6G). Taken altogether, these data indicate that EPS15 is a key component of Wnt9a/Fzd9b endocytosis and signaling, and that both domains previously described to impact on EGFR endocytosis are required for this function (Fig. 7).

**Figure 7:**
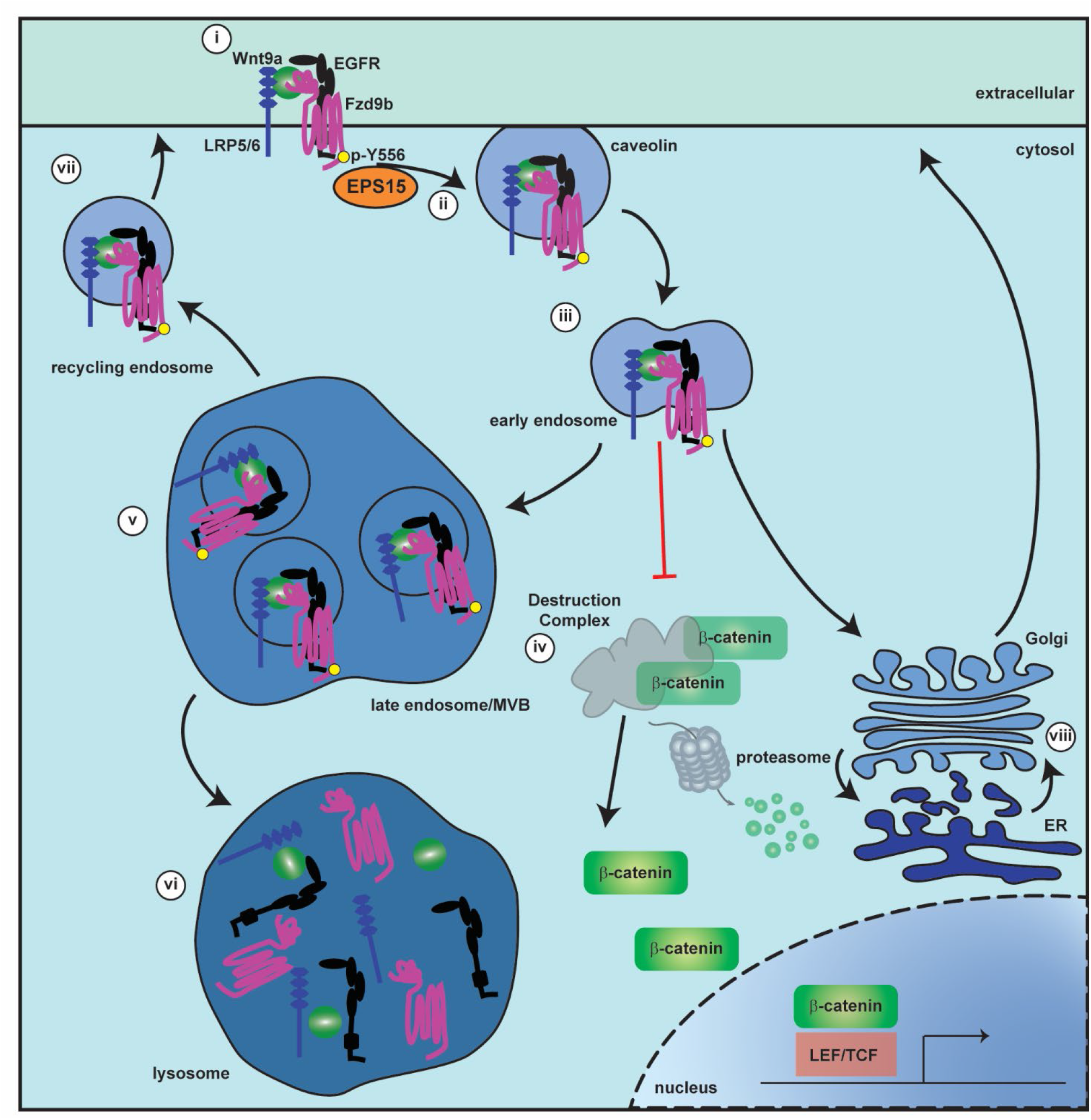
A model for Wnt9a/Fzd9b endocytosis, trafficking and signaling. **i.** When Wnt9a binds to the cell surface receptor complex including Fzd9b, EGFR and LRP5/6, EGFR phosphorylates the Fzd9b tail at Y556. **ii.** EPS15 and Caveolin are required for endocytosis of the complex into early endosomes (**iii**), where the receptor complex inhibits the destruction complex, allowing β-catenin release and translocation to the nucleus for target gene expression (**iv**). The Wnt9a/Fzd9b complex is further trafficked to the late endosome/multi-vesicular body (MVB, **v**), the lysosome (**vi**), the recycling endosome (**vii**), and the endoplasmic reticulum (ER, **viii**).

## Discussion

The Wnt pathway regulates a plethora of downstream biological events, likely through a medley of diverse transcriptional and non-transcriptional outputs driving changes in a panoply of cell types and tissues. At least part of the complexity of this system lies in the multitude of ligands (at least 19 in vertebrates) and receptors (at least 10 in vertebrates). One of the bigger mysteries in Wnt biology is deciphering how the different Wnt ligands choose their cognate receptors and co-receptors, and how these lead to the diverse outputs seen in development, homeostasis, and disease. One way to begin deciphering these mechanisms of specific Wnt/Fzd interactions is to visualize cell biological events involving known cognate pairings, which we have done here using the pairing of Wnt9a and Fzd9b, we previously demonstrated are required for HSPC proliferation in developing zebrafish and human cells^8, 9^.

The ability to monitor functional Wnt ligands has been challenging since Wnt fusion proteins often lose functionality. There are a small handful of examples of successfully tagging Wnt proteins with fluorophores; these have proven to be valuable in understanding cell signaling biology. For example, GFP-tagged mouse Wnt3a was used to visualize differences in binding of the promiscuous Wnt3a ligand to different Fzd proteins^46^, as well as to monitor how diffusion of Wnt3a is regulated^47^. *Xenopus* wnt2b-GFP provided insights into how Wnts can move along cellular protrusions^48^. Chick WNT1-GFP was used to show how Wntless is involved in changing the distribution of WNT1^49^. Imaging *Xenopus* wnt5a-GFP suggests receptor clustering at the membrane is important for signaling^50^, and zebrafish GFP-Wnt8 can be seen traveling from one cell to another in early neural plate development^51^. The field has therefore only begun to exploit visualizing Wnt ligands.

We demonstrate here that the location of the tag, as well as the fluorophore and linker sequence are all important to consider when deriving Wnt fusion proteins. Interestingly, mouse Wnt3a and *xenopus* Wnt8 tolerate GFP tagging at the N-terminus^46, 47, 51^, while we found that C-terminal tagging of Wnt9a was better tolerated (Fig. S2C), as has also been reported for *Xenopus* wnt2b^48^, chick WNT1^48^ and^48^, *Xenopus* Wnt5a^50^. These differences may be because of differences in location, or linker sequence and/or length, and they likely need to be determined empirically for each Wnt.

By administering our biologically functional Wnt9a-Gam protein to Fzd9b-mKate cells, we were able to visualize endocytosis and trafficking of the Wnt9a/Fzd9b (Fig. 3). Our data indicate that this endocytosis destines the complex for either degradation or recycling (Fig. 7). Non-specific small molecule inhibition of endocytosis leads to a loss of the nuclear β-catenin mediated Wnt signal (Fig. S3B-C), which also occurs in response to loss of CAV1 (Fig. 4B). These data are very interesting in the landscape of previous data from the Wnt field. For instance, neither clathrin, nor caveolin-mediated endocytosis are required for Wnt3a signaling in mouse embryonic stem cells^38^. However, a previous report from the same group suggests that mouse Wnt3a signaling is sensitive to some chemical inhibitors of endocytosis such as chlorpromazine, hypertonic sucrose or monodansylcadaverine^37^. As the authors suggest, the differences between the two systems examining mouse Wnt3a likely lies in the now well documented off-target effects of these chemical modulators^38, 70^. In HeLaS3 cells, the co-receptor LRP6 is internalized through caveolin in response to mouse Wnt3a, and this is required for STF activity; however, it is not clear if the Wnt3a ligand is also endocytosed^35^. In addition, further studies indicate that the observed differences in downstream signaling may actually be related to the importance of clathrin and AP2 for forming LRP6 signalosomes at the membrane^71^, or trafficking of the LRP6 receptor^72, 73^, rather than for endocytosis of the receptor complex itself. In contrast to mouse Wnt3a, the *Drosophila* ligand, Wingless (Wg), has been shown to require Rab5-mediated endocytosis^31^, and Vacuolar H+-ATPase-mediated acidification has been shown to be required for Wnt3a signaling^74^, suggesting that at least some Wnt ligands need to be internalized for function. Taken altogether, and in context of our findings, it stands to reason that one mechanism of the context-dependent specificity required downstream of different Wnt/Fzd pairings lies within endocytosis.

Mechanistically, we have demonstrated that Wnt9a/Fzd9b endocytosis requires EGFR-phosphorylation of the Fzd9b tail at Y556 (Fig. 5), and phosphorylation of Y850 of the adaptor protein EPS15 (Fig. 6G), as well as the N-terminal EH domains of EPS15 (Fig. 6G), to initiate caveolin-mediated endocytosis. The requirement for co-receptors in specific Wnt signals is starting to be better appreciated as a mechanism of regulating downstream signaling specificity^75–79^. These observations support the overall notion that co-receptors with enzymatic activity may be recruited to various Wnt/Fzd pairings, resulting in different types of endocytosis. These endocytic events may then either activate, or inhibit the Wnt signal, likely depending on the cellular and molecular context.

## Methods

### Cell culture

We had previously established HEK293 cells with a stably integrated Super-TOP-Flash reporter (STF)^45^, which were used for all luciferase assays, or to further derive stably integrated transgenic lines. HEK293, HEK293T, and Chinese hamster ovary (CHO) cells were grown in Dulbecco’s modified Eagle’s medium (DMEM) supplemented with 10% heat-inactivated fetal bovine serum (FBS) and 1% penicillin/streptomycin under standard conditions. Cells were routinely tested for mycoplasma contamination, which was always negative.

### Generation of transgenic and knockout cell lines and conditioned medium

LRP6 knockout HEK293 STF cells were previously generated^8^; LRP5/6 double knockout (DKO) cells were generated by transfecting a confluent 100mm plate of LRP6 knockout HEK293T STF cells with 3mg each of Cas9^80^ and two guide RNAs under regulatory control of a U6 promoter (GGAAAACTGGAAGTCCACTG and GCAGGACCTTGACCAGCCGA) targeting beta-propeller 1 of LRP5. Single cell clones were validated for loss of LRP5 coding sequence by sequencing the genomic locus using primers TCGCCGCTCCTGCTATTTG and AGGTCGATGGTCAGTCCATT, and immunoblotting using a rabbit monoclonal antibody (D80F2, Cell Signaling Technologies, 5731S). EGFR knockout cells were generated in a similar fashion using two guide RNAs (TAACAAGCTCACGCAGT and GGTGGTCCTTGGGAATT), which target early coding sequences in the second exon. Single cell clones were validated by sequencing the genomic locus with primers CTGCTACCCTTAATACCTGGAC and CCAGGCCTTTCTCCACTTAG, and immunoblotting with rabbit monoclonal antibody (EP38Y, Abcam, ab52894). EGFR knockout cells were also validated by testing for loss of tyrosine phosphorylation in response to a one-minute treatment with 50 ng/mL mouse EGF by immunoblotting with mouse monoclonal antibody (PY20, BioLegend, 309301).

Transgenic cell lines were generated using the PiggyBAC transposase system^81^. Briefly, 100mm plates of parental cell lines were transfected using polyethyleneimine (PEI), with 5ug5 ug of hyperactive PiggyBAC transposase, and 5 ug of plasmid harboring the cargo (including antibiotic resistance cassettes) flanked by PiggyBAC recombination sites. Individual clonal lines were generated using antibiotic selection and standard methods. Clonal lines were validated using immunoblotting and luciferase reporter assays, compared with untagged sequences.

Stable transgenic CHO cell lines harboring doxycycline-inducible Wnt proteins were grown to confluence in 150 mm plates, supplemented with 500 ng/mL doxycycline, and cultured for 10 days further, with additional doxycycline spiked in at 500 ng/mL every 3 days. Conditioned medium was collected from parental (mock CM), or transgenic (Wnt CM) lines, filtered sterilized with a 0.22 µm filter and validated for Wnt activity using STF assays. CM was considered active if Wnt CM was at least 40X more active than mock CM. Active CM was stored aliquoted at −80°C and diluted 1:2 with fresh medium prior to use on cells.

### Luciferase reporter STF assays

Luciferase reporter assays were performed with either transient transfection, or stable lines, as indicated in the figure legends. For transient transfection, 293 STF cells were seeded into twelve-well plates and transfected using polyethyleneimine (PEI), 50 ng of *renilla* reporter vector, 200 ng of Wnt or Fzd expression vector, with a total of 1 ug of DNA/well. When stable lines and conditioned medium were used, cells were seeded into twelve-well plates and after 24 hours were treated with a 1:1 mixture of fresh and conditioned medium. For siRNA experiments, a 6 well plate was treated with 30 pmol of siRNA using RNAiMax transfection reagent (Invitrogen) according to the manufacturer’s recommendations. All transfected cells were harvested 48 hours post-transfection and all conditioned medium or co-cultured cells were harvested 24 hours post-treatment; the lysates processed and analyzed using the Promega Dual Luciferase Assay System according to the manufacturer’s instructions. Each experiment was performed with at minimum biological triplicate samples and reproduced at least once with a similar trend. Wnt activity was calculated by normalizing Firefly Luciferase output to Renilla Luciferase; Wnt9a/Fzd9b fold induction was set to 100.

### Wnt protein enrichment and immunoblotting

Enrichment of Wnt proteins was accomplished using a Blue Sepharose pulldown, based on previously established Wnt purification methods^82^. Briefly, 2.5 mL of CM was supplemented with 1% Triton X-100, 20 mM Tris-Cl, and 0.1% NaN_3_, and filtered through a 0.22 µm syringe filter. Blue Sepharose 6 Fast Flow Agarose beads (GE, 17-0948-01) were washed with PBS and resuspended in a 50:50 PBS:beads slurry. A mixture of 2 mL of Wnt CM and 20 µL of Blue Sepharose were incubated for 90 minutes, rocking end over end at room temperature. Beads were collected by centrifugation at 400 g for 5 minutes and washed three times with 1 mL cold Wnt wash buffer (1% CHAPS, 150 mM KCl, 50 mM Tris-Cl in PBS). Wnt proteins were dissociated from the beads by heating to 95°C in Laemmli buffer^83^ for 5 minutes.

For cell lysate immunoblots, cells were harvested 48 hours after transfection in TNT buffer (1% Triton X-100, 150 mM NaCl, 50 mM Tris, pH8.0), with protease inhibitors. Immunoblots were performed according to standard procedures (lysates subjected to blotting for Fzds were not boiled), using antibodies for: α-tRFP, [1:1,000, mKate antibody, Evrogen, AB233], α-FLAG, [1:1,000, Sigma, F1804], α-V5, [1:5,000, Cell Signaling Technologies, 13202S], α−β−actin, [1:10,000, Sigma, A2228], 1:500, α-Fzd9b^8^, 1:500, α-Wnt9a^10^, α-CAV1, [Cell Signaling Technologies, 3267], α-Clathrin, [Invitrogen, MA1-065], α-EPS15, [R&D Systems, AF8480], α-LRP6, [Cell Signaling Technologies, 2560S], α-LRP5, [Cell Signaling Technologies, 5731S], α-EGFR, [Abcam, ab52894], α-pTyr, [BioLegend, 309301], α-β-catenin (total) [Cell Signaling Technologies, 8480S], α-activated-β-catenin [Cell Signaling Technologies, 8814S], α-rabbit-HRP, [1:20,000, Southern Biotech, 4050-05], α-mouse-HRP, [1:20,000, Southern Biotech, 1013-05].

### Plasmids

Expression constructs for fusion proteins were generated by standard means using PCR from plasmids harboring a CMV promoter, or a doxycycline inducible promoter system. Addgene provided expression vectors for *Cas9* (47929) and guide RNAs (46759).

### Confocal imaging and analysis

Glass coverslips (#1.5, 12mm diameter) were coated with Cultrex Poly-L-Lysine (R&D systems, 34-382-0001) for 1 hour at 37°C, washed 3 times with water and air dried. Cells were plated on coated coverslips for 24 hours prior to addition of CM in an equal volume to the culture medium already present. Cells were fixed by removing medium, washing three times with ice cold PBS^++^ (containing Mg^++^ and Ca^++^) and incubating with 4% paraformaldehyde (PFA) for 20 minutes at room temperature. For direct imaging of fluorescently labeled transgenic cells, slides were mounted using vectashield mounting medium and imaged within one week.

For immunofluorescence, cells were prepared and fixed as above, and incubated in blocking buffer (2% BSA in PBS^++^) for 60 minutes at room temperature, and in primary antibody (α-tRFP, [mKate antibody, Evrogen, AB233], α-FLAG, [Sigma, F1804], α-V5, [Cell Signaling Technologies, 13202S], α-Fzd9b^8^, α-Wnt9a^10^, α-Rab5, [Cell Signaling Technologies, 3547S], α-Rab7, [Cell Signaling Technologies, 9367S], α-Rab9a, [Proteintech, 11420-1-AP], α-Gpr94, [Cell Signaling Technologies, 20292], α-LAMP1, [Invitrogen, MA5-29385]) diluted 1:50-1:250 in blocking buffer overnight at 4°C. Cells were washed three times in PBS^++^, incubated in secondary antibody (1:500 for each, Goat-α-mouse Alexa 488 [cat# A11001, lot# 2189178], goat-α-rabbit Alexa 488 [cat# 4050-30, lot# e1317 NC39], goat-α-mouse Alexa 555 [cat# 1030-32, lot# E2518 ZA08E], goat-α-rabbit Alexa 647 [cat# 4050-31, lot# E1519 S109Z], Southern Biotech) diluted in blocking buffer for 60 minutes at room temperature, washed three times in PBS^++^, and mounted in vectashield mounting medium (with or without DAPI). Transferrin-488 internalization was performed essentially as previously described ^38^, 48 hours post siRNA transfection. Slides were imaged within one week.

Confocal z-slices were collected using a Zeiss LSM 880 with Airyscan, equipped with 405, 458, 488, 514, 561 and 633 nm laser lines and 5x air, 10x air, 20x air, 40x water, 63x oil and 100x oil objectives. Scanning was sequential and images were collected at a minimum of 1900 x 1900 resolution; images were processed using Airyscan Fast mode. Representative images were produced by combining 1-10 z-slices. Calculation of % events was performed by examining z-stacks through the entire cell and identifying cells that had internalized green puncta and dividing by the total number of cells in the field of view.

### Zebrafish husbandry and in situ hybridization

Zebrafish were maintained and propagated according to Van Andel Institute and local Institutional Animal Care and Use Committee policies. AB* zebrafish were used in all experiments. The MO for *eps15* was targeted to block the ATG start codon with sequence 5’-TCAGGGAGAGACTGGCAGCCAT −3’ from GeneTools. 1-cell stage zygotes were injected with 0.5ng of *eps15* MO. Rescue experiments were performed using 100 pg of Wnt9a-Gamillus expressing plasmid.

RNA probe synthesis was carried out according to the manufacturer’s recommendations using the DIG-RNA labeling kit, or the fluorescein labeling kit for FISH (Roche). Probe for *cmyb* and WISH protocols have been previously described^84–86^.

## Acknowledgements

The authors thank Hannah Morris-Little and Jordan Setayesh for technical assistance in generating EGFR and LRP5/6 DKO cells, respectively. Imaging, analysis, and quantification was performed in part by the Van Andel Institute Optical Imaging Core. Research reported in this publication was supported by the National Heart, Lung, And Blood Institute of the National Institutes of Health under Award Number R00HL133458, and by the National Institute of General Medical Science under Award Number R35GM142779. The content is solely the responsibility of the authors and does not necessarily represent the official views of the National Institutes of Health.

## Author Contributions

NN, KAC, KET, EM and CG designed and conducted experiments and analysis and edited the manuscript. SG conceived, designed, and supervised experiments and analysis, and wrote the manuscript.

## Correspondence

Correspondence and requests for materials should be addressed to SG at.

## Supplementary Information

**Supplementary Figure 1:**
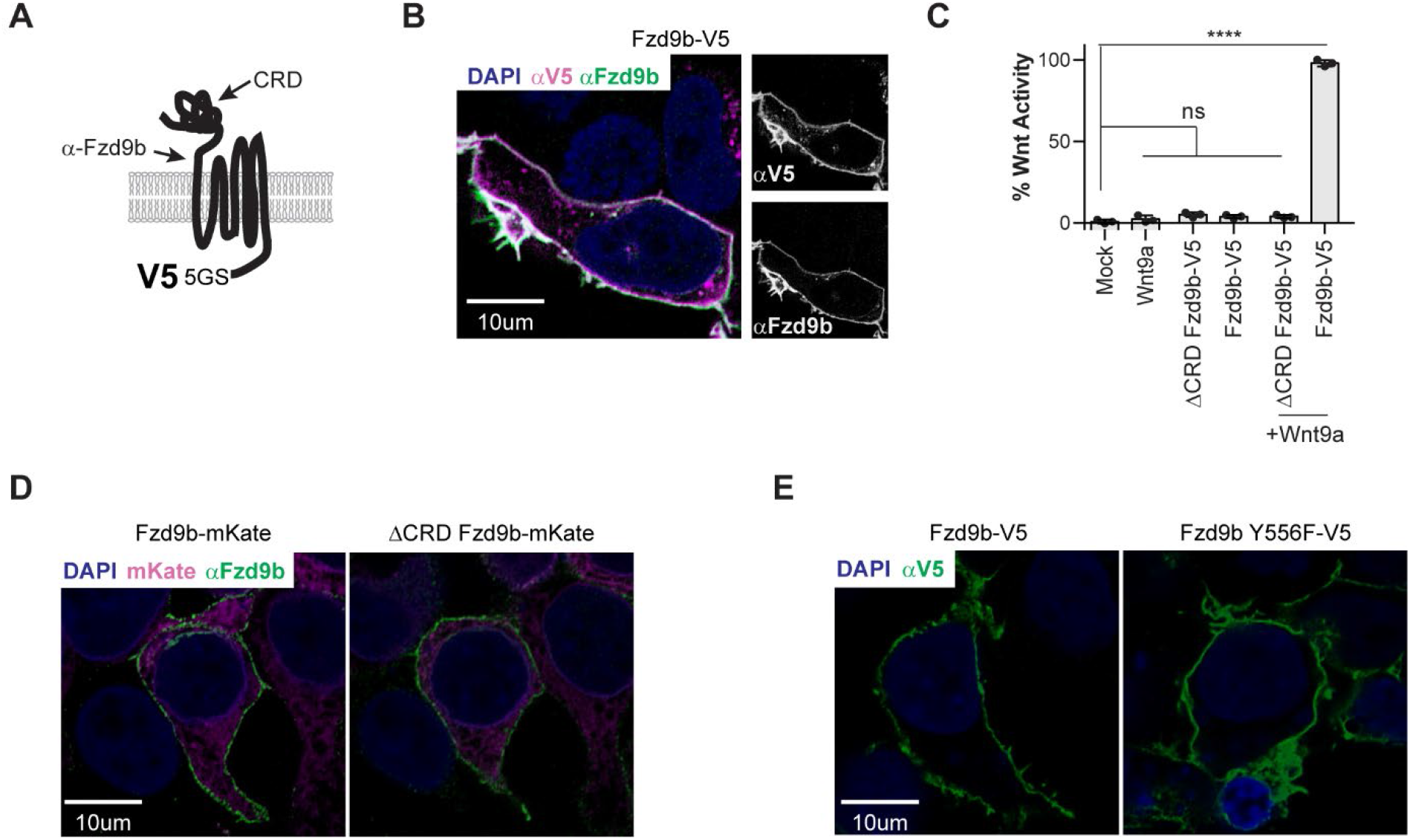
V5-tagged Fzd receptors maintain signaling capacity. **A.** Schematic of Fzd9b-V5. **B.** Confocal z-stacks of Fzd9b-V5 cells with immunofluorescence for Fzd9b (green) and V5 (magenta), DAPI (blue). **C**. STF assays for Fzd-V5 fusion proteins treated with Wnt9a CM. **D.** Confocal z-stacks of Fzd9b-mKate and ΔCRD Fzd9b-mKate cells with non-permeabilized immunofluorescence for Fzd9b (green); mKate (magenta), DAPI (blue). **E.** Confocal z-stacks of Fzd9b-V5 and Fzd9b Y556F-V5 cells with non-permeabilized immunofluorescence for Fzd9b (green); blue is DAPI (nuclei). n.s. not significant, ****P<0.0001 by ANOVA with Tukey post-hoc comparison.

**Supplementary Figure 2:**
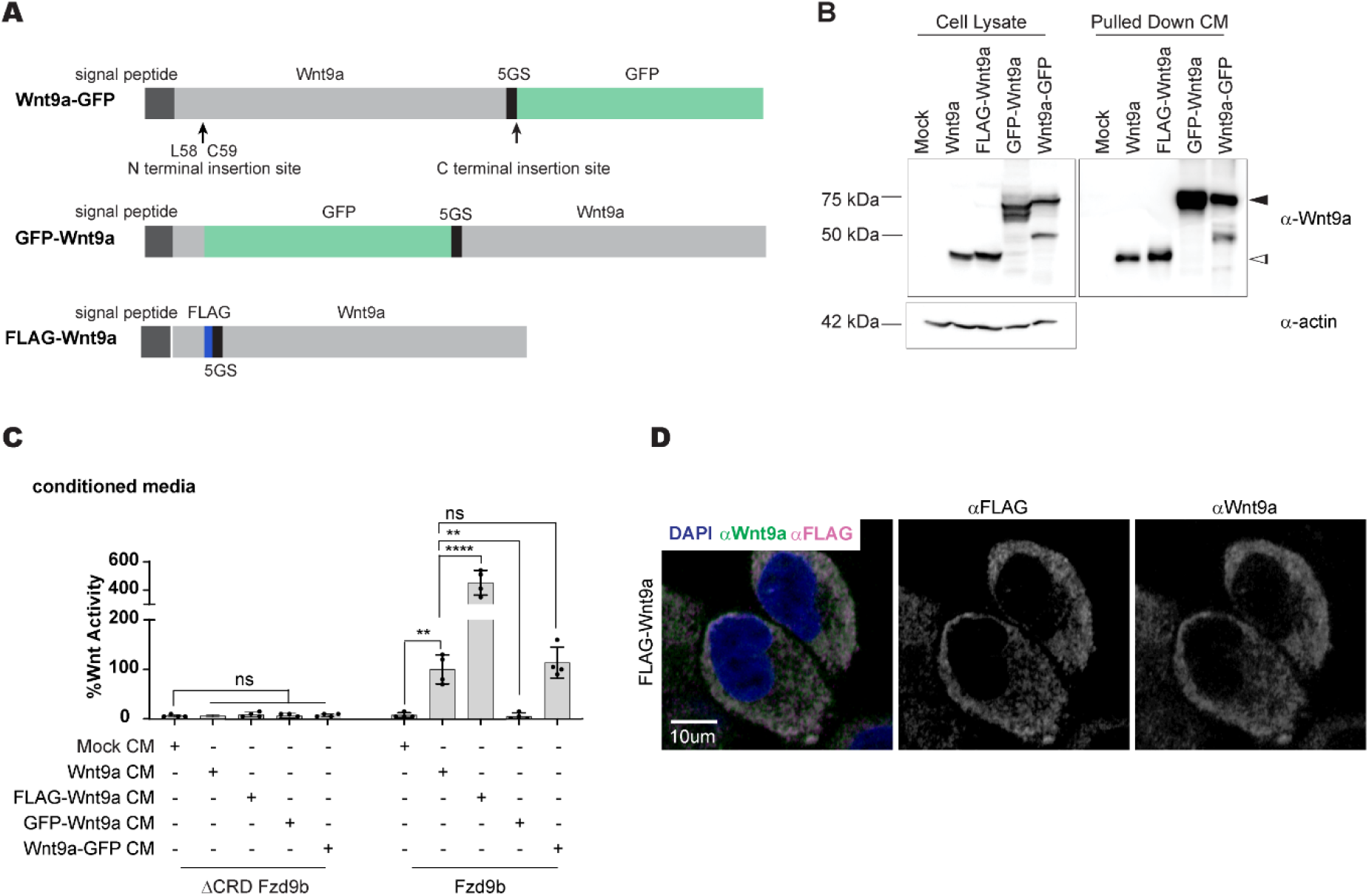
Tagging Wnt9a impacts on signaling capacity. **A.** Schematics of Wnt9a fusion proteins. **B.** Immunoblots from cell lysates or enriched CM, blotted for Wnt9a and β-Actin from transgenic CHO cell lines expressing Wnt9a constructs as shown. Black arrowheads denote anticipated sizes of fusion proteins; white arrowheads denote anticipated size of WT Wnt9a. **C**. STF assays in Fzd9b-mKate STF cells, treated with Wnt9a fusion protein CM. **D.** Confocal z-stacks of FLAG-Wnt9a CHO cell immunofluorescence for Wnt9a (green) and FLAG (magenta), DAPI (blue). n.s. not significant, **P<0.01, ****P<0.0001 by ANOVA with Tukey post-hoc comparison.

**Supplementary Figure 3:**
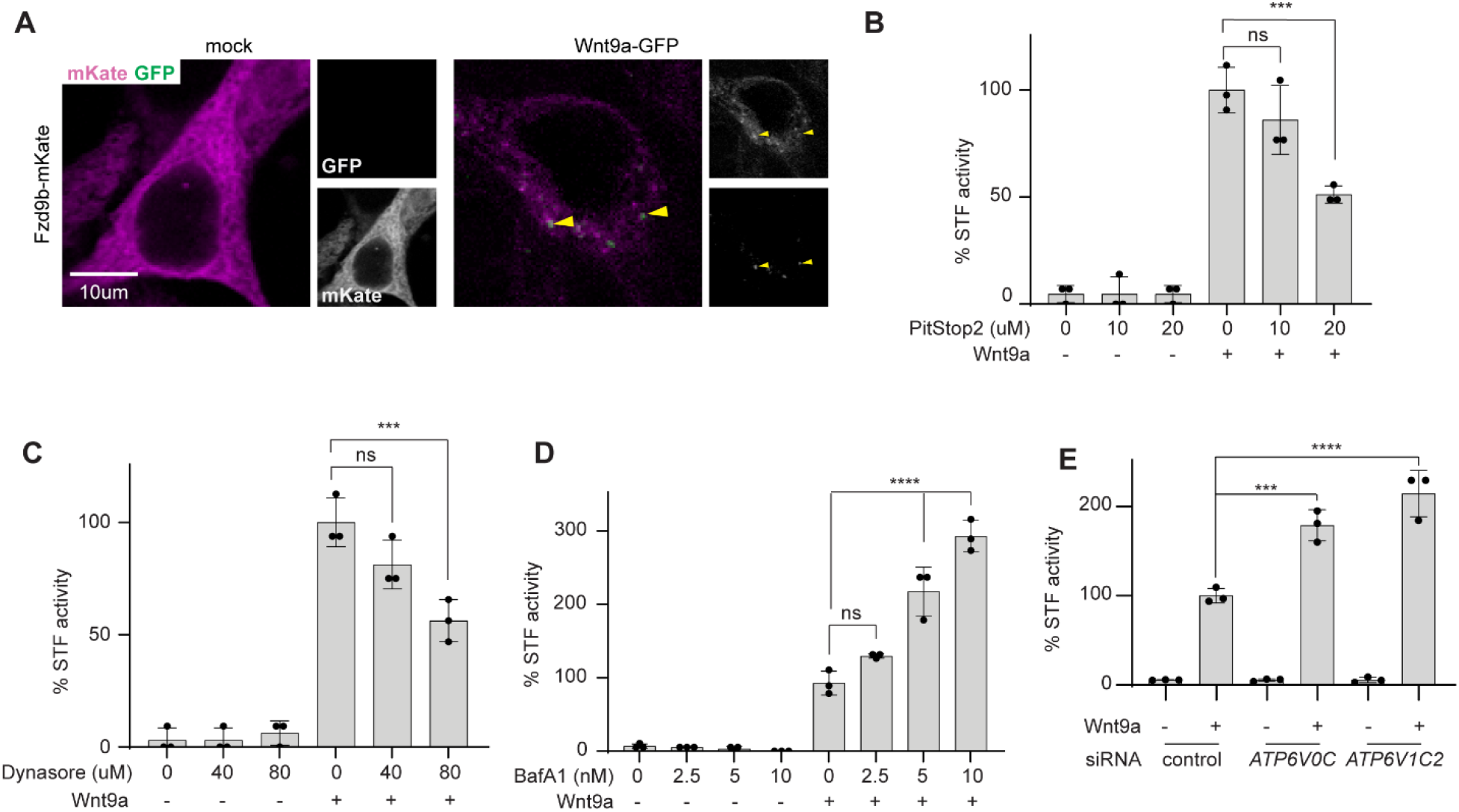
The Wnt9a/ Fzd9b complex is rapidly endocytosed. **A.** Confocal z-stacks of Fzd9b-mKate cells treated with either mock or Wnt9a-GFP CM and fixed at one minute. Yellow arrows indicate Wnt9a/Fzd9b complexes. Green is GFP and magenta is mKate. STF assays in Fzd9b-mKate STF cells, treated with Wnt9a CM in the context of PitStop2 (**B**), Dynasore (**C**), or BafilomycinA1 (**D**) at indicated dosages. **E.** STF assays in Fzd9b-mKate STF cells, treated with control, *ATP6V0C*, or *ARP6V1C2* siRNAs, and induced with Wnt9a CM. n.s. not significant, ***P<0.001, ****P<0.0001 by ANOVA with Tukey post-hoc comparison.

**Supplementary Figure 4:**
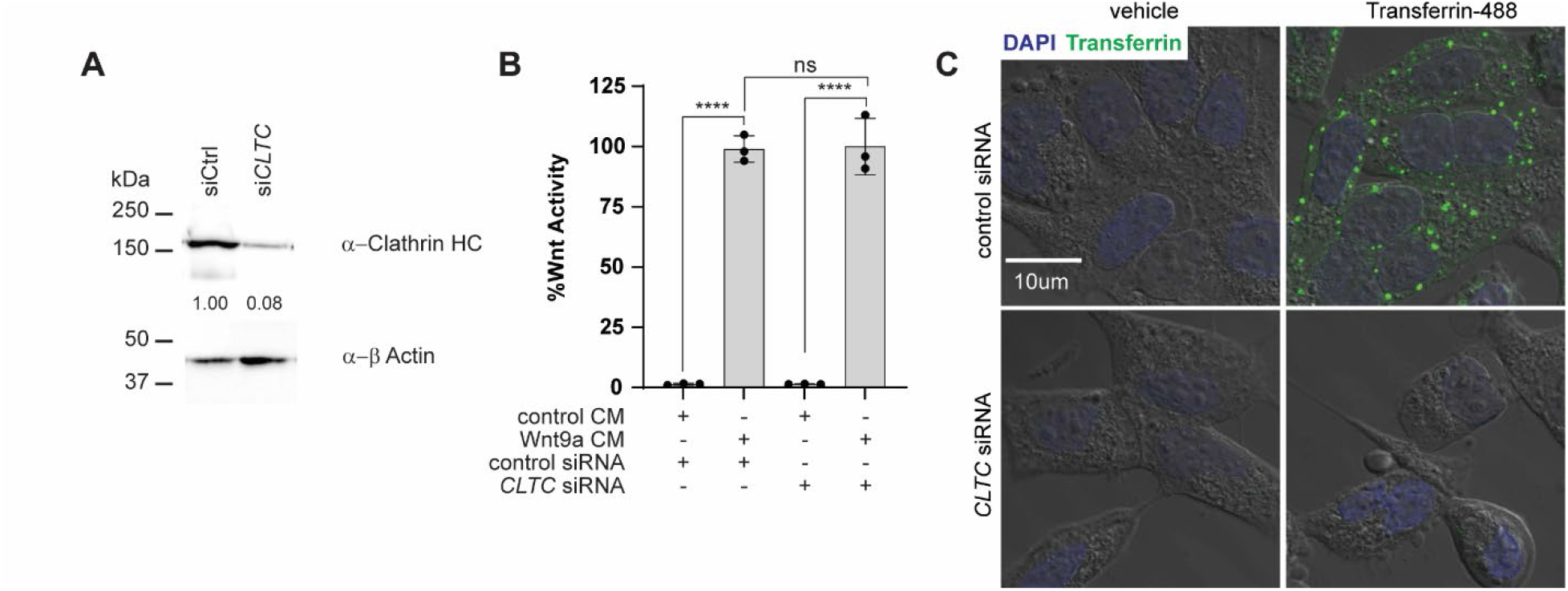
The Wnt9a/Fzd9b signal does not require Clathrin-mediated endocytosis for signaling. **A.** Immunoblots from cells treated with either control or *CLTC* siRNAs, blotted for Clathrin and β-Actin. **B**. STF assays in Fzd9b-mKate STF cells, treated with control or *CLTC* siRNAs, and induced with Wnt9a CM. **C**. single confocal Z-slice through HEK293T cells treated with either control of CLTC siRNAs and vehicle or Transferrin-488; DAPI (blue), transferrin (green) are overlaid on transmitted light image. n.s. not significant, ****P<0.0001 by ANOVA with Tukey post-hoc comparison.

**Supplementary Figure 5:**
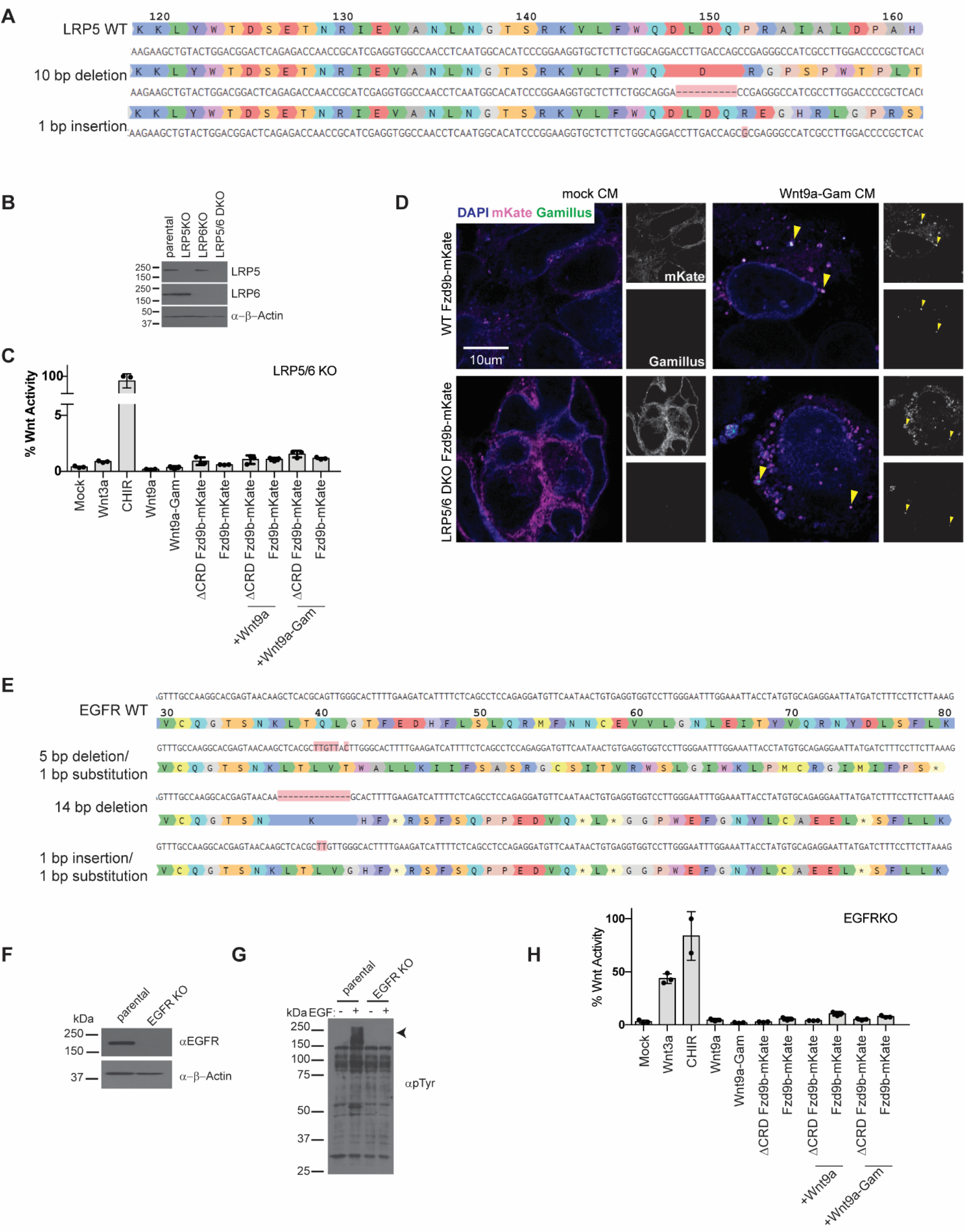
Validation of LRP5/6 and EGFR knockout lines. **A.** Sequences alignment of LRP5 WT (top) with 2 mutant alleles detected by NGS sequencing. Amino acid residues are numbered relative to start codon. **B.** Immunoblots from parental, LRP5 KO, LRP6 KO or LRP5/6 DKO cells, blotted for LRP5, LRP6 and β-Actin. **C**. STF assays in DCRD Fzd9b-mKate of Fzd9b-mKate LRP5/6 DKO STF cells, induced with mock, Wnt3a, Wnt9a, or Wnt9a-Gam CM, or the Wnt agonist CHIR. **D**. Confocal z-stacks of Fzd9b-mKate WT or LRP5/6 DKO cells treated with either mock or Wnt9a-GFP CM and fixed at one minute. Yellow arrows indicate Wnt9a/Fzd9b complexes. Green is GFP and magenta is mKate. **E.** Sequences alignment of EGFR WT (top) with 3 mutant alleles detected by NGS sequencing. Amino acid residues are numbered relative to start codon. **F.** Immunoblots from parental or EGFR KO cells, blotted for EGFR and β-Actin. **G**. Immunoblots from parental or EGFR KO cells, induced with 1nM EGF and blotted for p-Tyrosine. Arrowhead indicates anticipated size of other ERBB family members anticipated to be phosphorylated in response to EGF with functional EGFR. **H**. STF assays in DCRD Fzd9b-mKate of Fzd9b-mKate EGFR STF cells, induced with mock, Wnt3a, Wnt9a, or Wnt9a-Gam CM, or the Wnt agonist CHIR.

**Supplementary Figure 6:**
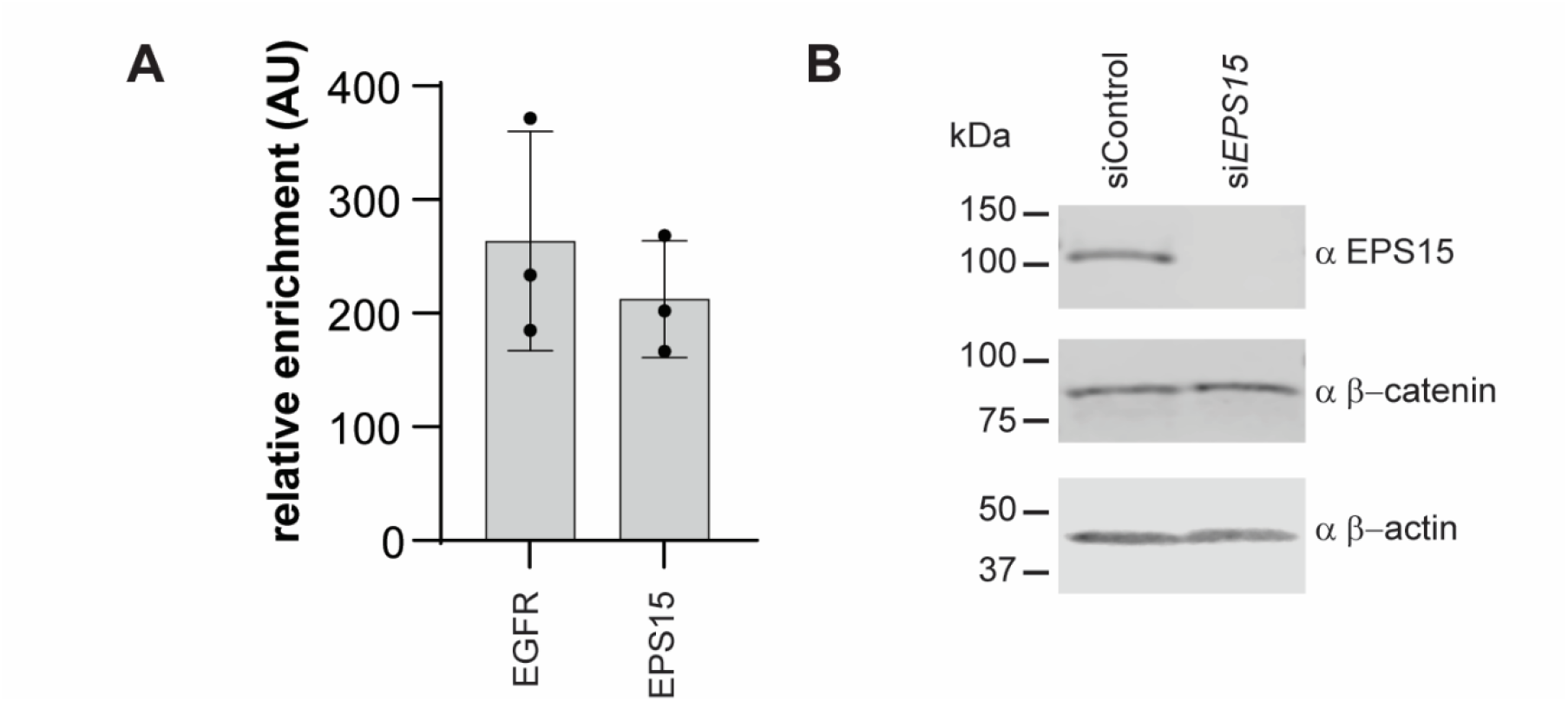
EPS15 is required for Wnt9a/Fzd9b endocytosis and signaling. **A.** Immunoblots from cell lysates of HEK293s treated with either control or *EPS15* siRNAs, blotted for EPS15 and β-Actin. **B.** Immunoblots from cells transfected with EPS15 WT and mutant construction, blotted for EPS15 and β-Actin.

