## Supplementary Data for "EGFR-initiated endocytosis of Wnt9a and Fzd9b is required for β-catenin signaling"

Nguyen, et al, 2022

Supplementary Information

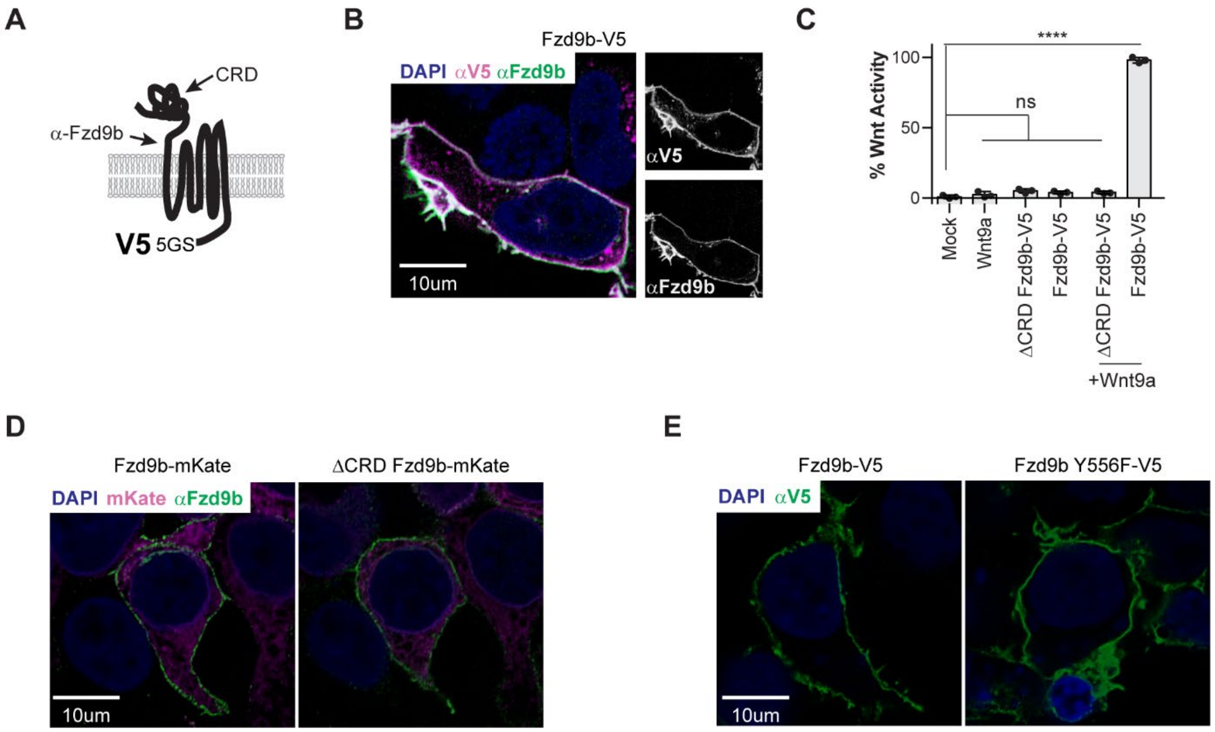


**Supplementary Figure 1: V5-tagged Fzd receptors maintain signaling capacity. A.** Schematic of Fzd9b-V5. **B.** Confocal z-stacks of Fzd9b-V5 cells with immunofluorescence for Fzd9b (green) and V5 (magenta), DAPI (blue). **C**. STF assays for Fzd-V5 fusion proteins treated with Wnt9a CM. **D.** Confocal z-stacks of Fzd9b-mKate and ΔCRD Fzd9b-mKate cells with non-permeabilized immunofluorescence for Fzd9b (green); mKate (magenta), DAPI (blue). **E.** Confocal z-stacks of Fzd9b-V5 and Fzd9b Y556F-V5 cells with non-permeabilized immunofluorescence for Fzd9b (green); blue is DAPI (nuclei). n.s. not significant, ****$P<0.0001$ by ANOVA with Tukey post-hoc comparison.

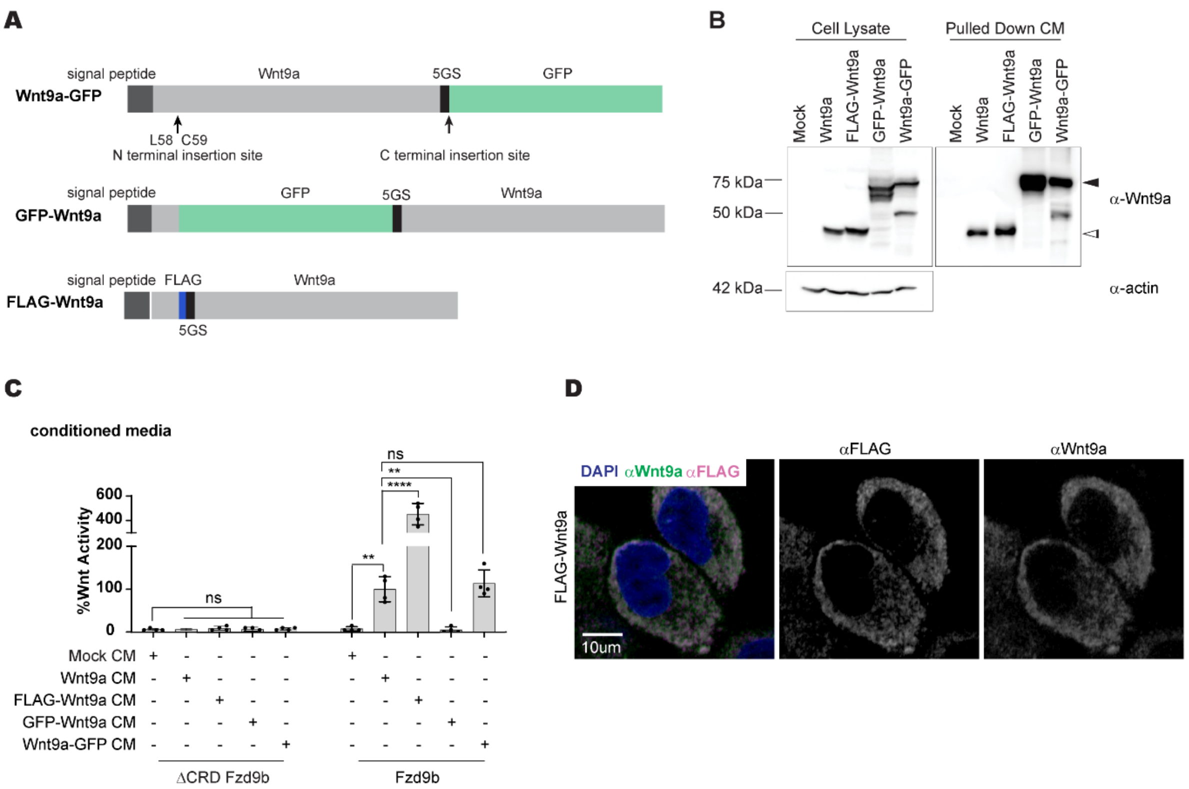


**Supplementary Figure 2: Tagging Wnt9a impacts on signaling capacity. A.** Schematics of Wnt9a fusion proteins. **B.** Immunoblots from cell lysates or enriched CM, blotted for Wnt9a and β-Actin from transgenic CHO cell lines expressing Wnt9a constructs as shown. Black arrowheads denote anticipated sizes of fusion proteins; white arrowheads denote anticipated size of WT Wnt9a. **C**. STF assays in Fzd9b-mKate STF cells, treated with Wnt9a fusion protein CM. **D.** Confocal z-stacks of FLAG-Wnt9a CHO cell immunofluorescence for Wnt9a (green) and FLAG (magenta), DAPI (blue). n.s. not significant, **P<0.01, ****P<0.0001 by ANOVA with Tukey post-hoc comparison.

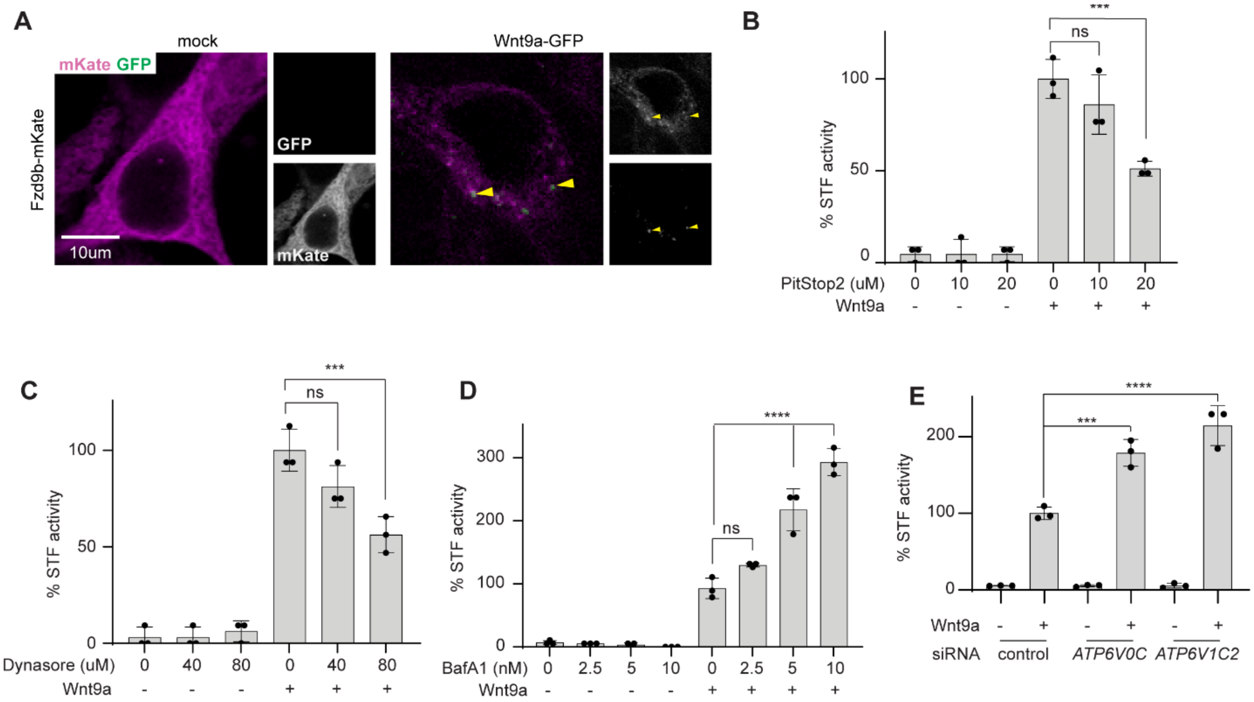


**Supplementary Figure 3: The Wnt9a/ Fzd9b complex is rapidly endocytosed. A.** Confocal z-stacks of Fzd9b-mKate cells treated with either mock or Wnt9a-GFP CM and fixed at one minute. Yellow arrows indicate Wnt9a/Fzd9b complexes. Green is GFP and magenta is mKate. STF assays in Fzd9b-mKate STF cells, treated with Wnt9a CM in the context of PitStop2 (**B**), Dynasore (**C**), or BafilomycinA1 (**D**) at indicated dosages. **E.** STF assays in Fzd9b-mKate STF cells, treated with control, *ATP6V0C*, or *ARP6V1C2* siRNAs, and induced with Wnt9a CM. n.s. not significant, ***P<0.001, ****P<0.0001 by ANOVA with Tukey post-hoc comparison.

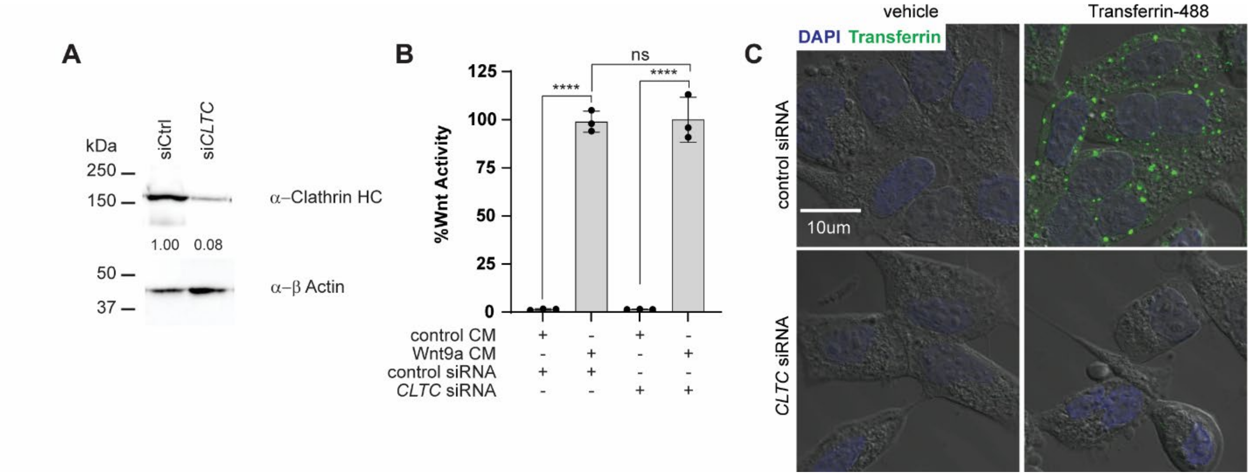


**Supplementary Figure 4: The Wnt9a/Fzd9b signal does not require Clathrin-mediated endocytosis for signaling. A.** Immunoblots from cells treated with either control or *CLTC* siRNAs, blotted for Clathrin and β-Actin. **B**. STF assays in Fzd9b-mKate STF cells, treated with control or *CLTC* siRNAs, and induced with Wnt9a CM. **C**. single confocal Z-slice through HEK293T cells treated with either control of CLTC siRNAs and vehicle or Transferrin-488; DAPI (blue), transferrin (green) are overlaid on transmitted light image. n.s. not significant, ****P<0.0001 by ANOVA with Tukey post-hoc comparison.

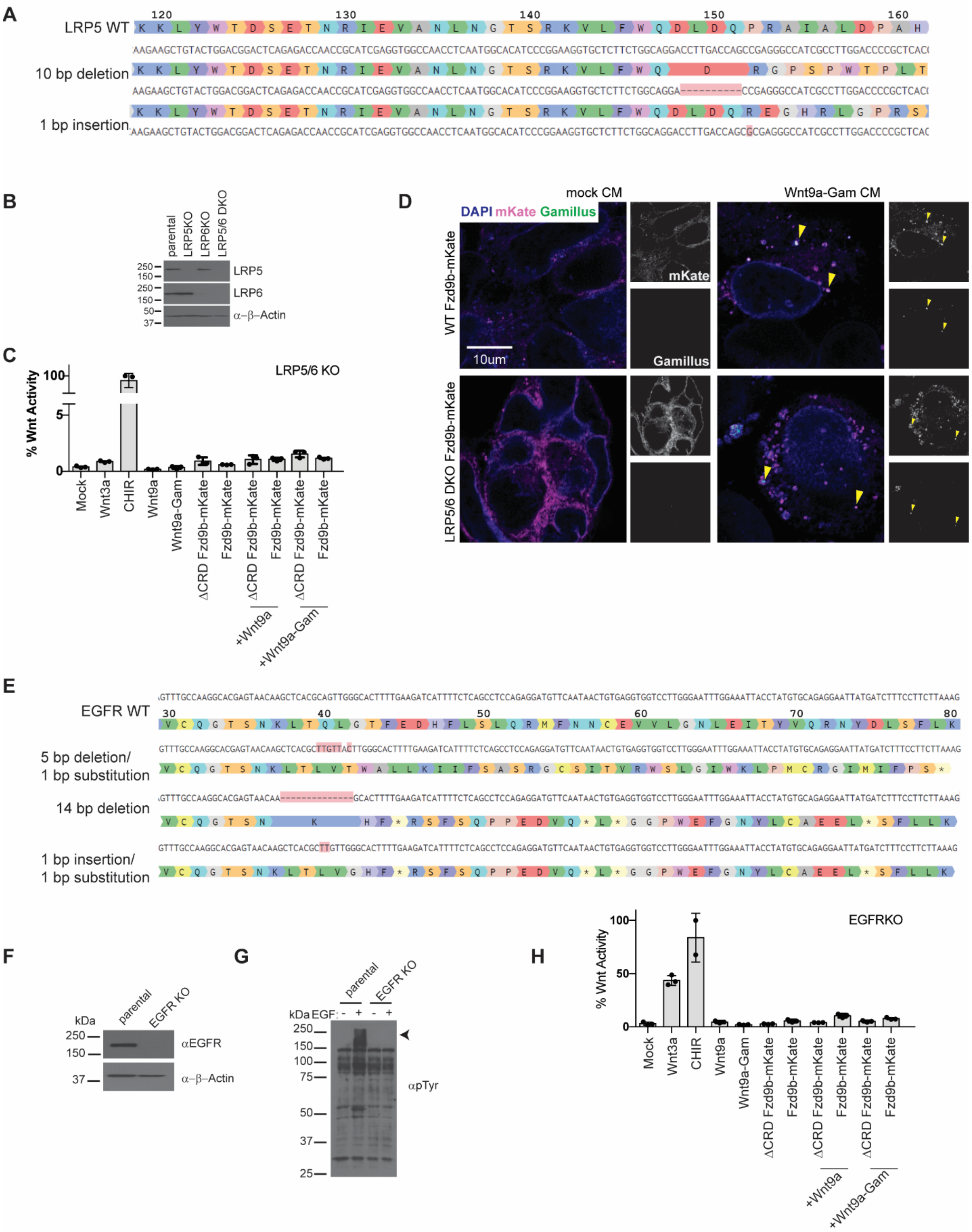


**Supplementary Figure 5: Validation of LRP5/6 and EGFR knockout lines. A.** Sequences alignment of LRP5 WT (top) with 2 mutant alleles detected by NGS sequencing. Amino acid residues are numbered relative to start codon. **B.** Immunoblots from parental, LRP5 KO, LRP6

KO or LRP5/6 DKO cells, blotted for LRP5, LRP6 and β-Actin. **C**. STF assays in DCRD Fzd9b-mKate of Fzd9b-mKate LRP5/6 DKO STF cells, induced with mock, Wnt3a, Wnt9a, or Wnt9a-Gam CM, or the Wnt agonist CHIR. **D**. Confocal z-stacks of Fzd9b-mKate WT or LRP5/6 DKO cells treated with either mock or Wnt9a-GFP CM and fixed at one minute. Yellow arrows indicate Wnt9a/Fzd9b complexes. Green is GFP and magenta is mKate. **E.** Sequences alignment of EGFR WT (top) with 3 mutant alleles detected by NGS sequencing. Amino acid residues are numbered relative to start codon. **F.** Immunoblots from parental or EGFR KO cells, blotted for EGFR and β-Actin. **G**. Immunoblots from parental or EGFR KO cells, induced with 1nM EGF and blotted for p-Tyrosine. Arrowhead indicates anticipated size of other ERBB family members anticipated to be phosphorylated in response to EGF with functional EGFR. **H**. STF assays in DCRD Fzd9b-mKate of Fzd9b-mKate EGFR STF cells, induced with mock, Wnt3a, Wnt9a, or Wnt9a-Gam CM, or the Wnt agonist CHIR.

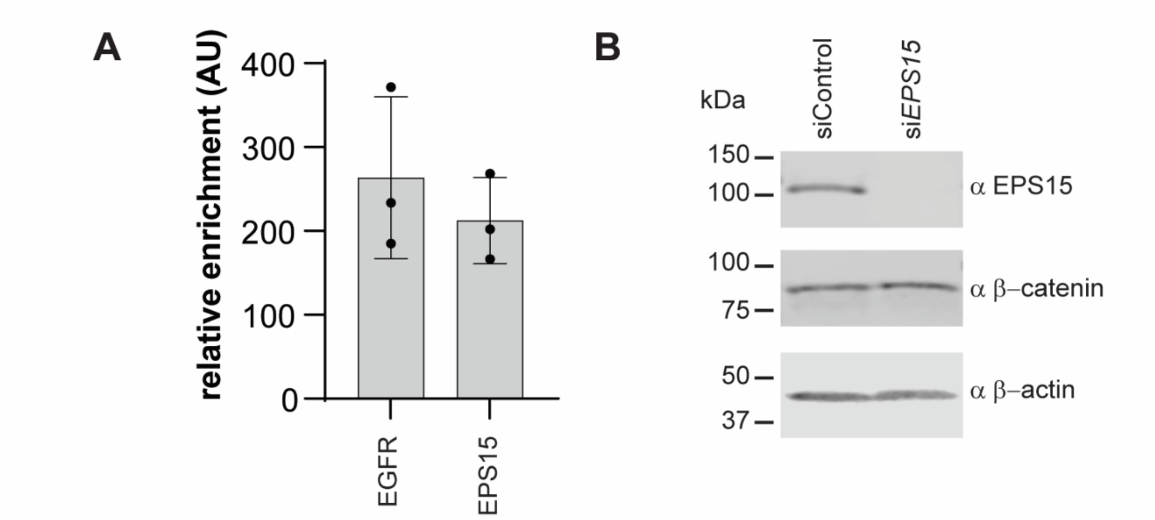


**Supplementary Figure 6: EPS15 is required for Wnt9a/Fzd9b endocytosis and signaling. A.** Immunoblots from cell lysates of HEK293s treated with either control or *EPS15* siRNAs, blotted for EPS15 and β-Actin. **B.** Immunoblots from cells transfected with EPS15 WT and mutant construction, blotted for EPS15 and β-Actin.
